# Poc1 is a basal body inner junction protein that promotes triplet microtubule integrity and interconnections

**DOI:** 10.1101/2023.11.17.567593

**Authors:** Marisa D. Ruehle, Sam Li, David A. Agard, Chad G. Pearson

**Affiliations:** Department of Cell and Developmental Biology, University of Colorado Anschutz Medical Campus, Aurora, CO, USA; Department of Biochemistry and Biophysics, University of California San Francisco, San Francisco, CA, USA; Chan Zuckerberg Institute for Advanced Biological Imaging, 3400 Bridge Parkway, Redwood Shores, CA, USA

**Keywords:** cilia, basal body, microtubule, inner junction Running title: Poc1 is a basal body inner junction protein

## Abstract

Basal bodies (BBs) are conserved eukaryotic structures that organize motile and primary cilia. The BB is comprised of nine, cylindrically arranged, triplet microtubules (TMTs) that are connected to each other by inter-TMT linkages which maintain BB structure. During ciliary beating, forces transmitted to the BB must be resisted to prevent BB disassembly. Poc1 is a conserved BB protein important for BBs to resist ciliary forces. To understand how Poc1 confers BB stability, we identified the precise position of Poc1 binding in the *Tetrahymena* BB and the effect of Poc1 loss on BB structure. Poc1 binds at the TMT inner junctions, stabilizing TMTs directly. From this location, Poc1 also stabilizes inter-TMT linkages throughout the BB, including the cartwheel pinhead and the inner scaffold. Moreover, we identify a molecular response to ciliary forces via a molecular remodeling of the inner scaffold, as determined by differences in Fam161A localization. Thus, while not essential for BB assembly, Poc1 promotes BB interconnections that establish an architecture competent to resist ciliary forces.

## INTRODUCTION

Basal bodies (BBs) and centrioles organize cilia and centrosomes, respectively, and are one of the most evolutionarily ancient structures of eukaryotic cells (Carvalho-Santos et al., 2011). They have diverse and essential functions in cell signaling, cell division, spermatogenesis, and development. These functions depend largely on the role of BBs in the nucleation and anchoring of cilia to the cell cortex (Carvalho-Santos et al., 2011; Wickstead and Gull, 2011). BBs are microtubule-based structures, comprised of nine radially arranged triplet microtubules (TMTs) creating a cylinder approximately 450 nm in length and 200 nm in diameter. TMTs consist of one full, 13-protofilament MT (the A-tubule), and two partial MTs (the B- and C-tubules), which are assembled on the exterior side of the previous tubule. The places where the tubules meet are referred to as junctions, and are further specified as inner junctions if they face the BB lumen, or as outer junctions if they are on the exterior of the BB. BBs dock to the cell cortex and the ciliary axoneme is formed from doublet MTs (DMTs) that extend continuously from the A- and B-tubules of the BB TMTs. The resulting cilia have unique functions depending on whether or not they are motile. Immotile cilia primarily operate as a signaling and environment sensing organelle. On the other hand, motile cilia contain axonemal dynein motor proteins that slide the cilia DMTs relative to each other and, when resisted by nexin links and the BB, create a beat stroke that is important for fluid movement and cell motility. When DMTs of the cilium are continuous with the TMTs of the BB, the forces generated by ciliary beating are transmitted directly to the BB (Dippell, 1968; Allen, 1969; Dirksen, 1971). Thus, BBs must resist these forces both to maintain their structural integrity, as well as to transmit those forces to the cell. This stability is thought to be achieved in two ways. First, BB TMTs are subject to myriad post-translational modifications that confer stabilizing properties to the MT lattice (Wloga et al., 2017). Second, BBs contain hundreds of proteins that are associated with the TMTs (Li et al., 2012; Kilburn et al., 2007; Le Guennec et al., 2020; Andersen et al., 2003; Keller et al., 2005). These proteins can stabilize the microtubule lattice of the TMT directly and / or connect neighboring TMTs, thereby promoting TMT organization and BB shape. How these molecules establish stabilizing structures, the identity of the full repertoire of these molecules, and how they reinforce the BB architecture remain poorly understood.

Structural studies show that BBs from a wide range of organisms share the same structural features, and these features highlight how the nine TMTs of BBs are assembled and supported. Although species-specific differences exist, most commonly, the proximal region of the BB (∼100 nm) harbors the cartwheel, which is comprised of stacks of nine Sas6 homodimers (Allen, 1969; Dippell, 1968; Nakazawa et al., 2007). Sas6 forms radial spokes that emanate from a central hub (Kitagawa et al., 2011; van Breugel et al., 2011). The cartwheel is thought to, at least in part, establish the nine-fold arrangement of TMTs (Nakazawa et al., 2007; Hilbert et al., 2016). Each cartwheel spoke terminates with a density called the pinhead that interacts with the A-tubule of each TMT. CEP135 / Bld10 is an important component of the pinhead (Hiraki et al., 2007; Lin et al., 2013; Guichard et al., 2017). A/C linkers also reside in the proximal region of the BB (∼150 nm), where they connect the C-tubule of one TMT to the A-tubule of its adjacent TMT (Allen, 1969; Li et al., 2012). To date, molecules that comprise the A/C linkers have not been identified. In the medial ∼300 nm of the BB, termed the core region, an inner scaffold attaches to and reinforces the nine TMTs, along with unique core region A/C linking structures (Le Guennec et al., 2020; Allen, 1969). Recent advances in Ultrastructure Expansion Microscopy (UExM) led to the identification of components of the inner scaffold in human centrioles, namely, Fam161A, Poc1B, Centrin2, Poc5, HAUS6, γ-tubulin, and CCDC15 (Arslanhan et al., 2023; Le Guennec et al., 2020; Schweizer et al., 2021). However, the precise localization of these proteins in the scaffold architecture and how they link to the TMTs is not well understood. WDR90/Poc16 is a core region-localized protein that is important for recruitment of proteins to the inner scaffold (Steib et al., 2020; Hamel et al., 2017). The combination of electron microscopy and super resolution light microscopy has thus proved to be a powerful means to dissect structural and compositional features of the BB, but the structure-function relationships that promote BB stability in the face of cilia generated forces is still lacking.

To understand how the BB architecture enables force resistance, we studied the conserved BB stability and disease-related protein, Poc1 (Pearson et al., 2009a; Li et al., 2021; Beck et al., 2014; Hua et al., 2023). Poc1 is necessary for centriole integrity and enables BBs to resist mechanical forces from beating cilia. Studies in the ciliate, *Tetrahymena thermophila*, revealed that in the absence of Poc1, BBs assemble the stereotypical nine-fold symmetric BB structure that appears normal, but cilia forces cause BBs to disassemble (Pearson et al., 2009a; Meehl et al., 2016). Therefore, Poc1 loss provides a unique opportunity to understand structural features of BBs that specifically promote BB stability. Identifying the precise localization of Poc1 within the BB structure is central to this goal. One of the human homologs of Poc1, human Poc1B, localizes to the BB core region and is proposed to be a component of the inner scaffold (Le Guennec et al., 2020). However, in *Tetrahymena*, a significant portion of Poc1 protein population localizes to the BB proximal end (Pearson et al., 2009a). Consistent with a proximal localization of Poc1, *poc1Δ* mutants in *Tetrahymena* partially disrupt A/C linkers, leading to the hypothesis that Poc1 is an integral component of the A/C linkers, or at least stabilizes them (Meehl et al., 2016; Li et al., 2019). Therefore, how and if Poc1 contributes to structural features unique to different BB domains is unclear.

Poc1 is a WD40 domain containing protein, comprised of a seven-bladed β-propeller at its N-terminus that forms a doughnut-shaped structure. WD40 domains commonly provide scaffolds for assembly of protein complexes where components are recruited at different faces of the doughnut (Xu and Min, 2011). In addition, Poc1 contains a short N-terminal extension before the WD40 domain (37 amino acids in *Tt*Poc1), and a longer, C-terminal extension after the WD40 domain (292 amino acids in *Tt*Poc1), which likely exit the WD40 motif from the same β-propeller blade (Hudson and Cooley, 2008). Although the WD40 domain alone is sufficient to rescue the BB loss phenotype in the *Tetrahymena poc1Δ* mutant, Poc1’s C-terminal extension is conserved, suggesting it plays important roles as well (Pearson et al., 2009a; Fourrage et al., 2010). Most of the C-terminal extension is predicted to be unstructured in an unbound, apo state, but the very end of the C-terminal extension contains a predicted helix-forming region that has coiled-coil forming propensity. This helical region in human Poc1B directly interacts with the Fam161A inner scaffold protein (Roosing et al., 2014). Thus, Poc1 may use its WD40 domain as well as its N- and C-terminal extensions for its BB functions.

To understand how BBs resist ciliary forces, we used cryogenic electron tomography (cryoET) and subtomogram averaging to solve the structure of the *Tetrahymena* BB at different longitudinal positions of the BB. Focused refinement of subregions of the TMT structure reached subnanometer resolution. This approach, using WT and poc1Δ BBs, enabled us to identify the location of Poc1 at the inner junctions of the TMTs, as well as to understand the role of Poc1 in the establishment of stabilizing structures of the BB. We propose that Poc1 seals microtubules together on the luminal side of the BB and stabilizes the cartwheel pinheads at the BB proximal end and the inner scaffold in the BB central core. This promotes TMT interconnections throughout most of the BB length, which are critical for the organelle’s shape maintenance and force resistance. Together, these findings identify the TMT inner junctions as a key location from which multiple BB stabilizing strategies are organized.

## RESULTS

### Conserved ultrastructure of the *Tetrahymena* BB

To identify the 3D high-resolution structure of *Tetrahymena* BBs using cryoET, we developed a procedure to gently isolate BBs while preserving overall BB structure (Fig. 1A-B; Video 1). Subtomogram averages were generated on TMTs from the BB proximal 150 nm, the central core 300 nm, and the distal 100 nm at 19-27 Å resolution (Fig. 1C-E, Fig. S1, Table 1). As in other organisms, the proximal region contained pinhead densities attached to the A-tubules (protofilaments A03-A04), as well as A/C linkers between the A- and C-tubules (protofilaments A06-08, and C07-10, Fig. 1C, Fig. S2A). The core region contained an inner scaffold (Fig. 1D, Fig. S2B). Finally, the subtomogram average from the BB distal region lacked these microtubule interconnecting structures and consisted of only DMTs. Some loss of the C-tubule was observed in the subtomogram average in the core region and at the distal end of the BB. While nearly all TMTs in the proximal region were complete TMTs, the core region showed only 9.3% complete TMTs. Given that TMTs are apparent in the core region in thin section EMs of *Tetrahymena* BBs, the C-tubule loss likely occurred as a result of the isolation and/or freezing process (Giddings et al., 2010; Allen, 1969). This suggests that the core region C-tubule is less stable than the A- and B-tubules. In summary, the 3D domain architecture of *Tetrahymena* BBs is comparable to other organisms (Fig. S2).

**Figure 1:**
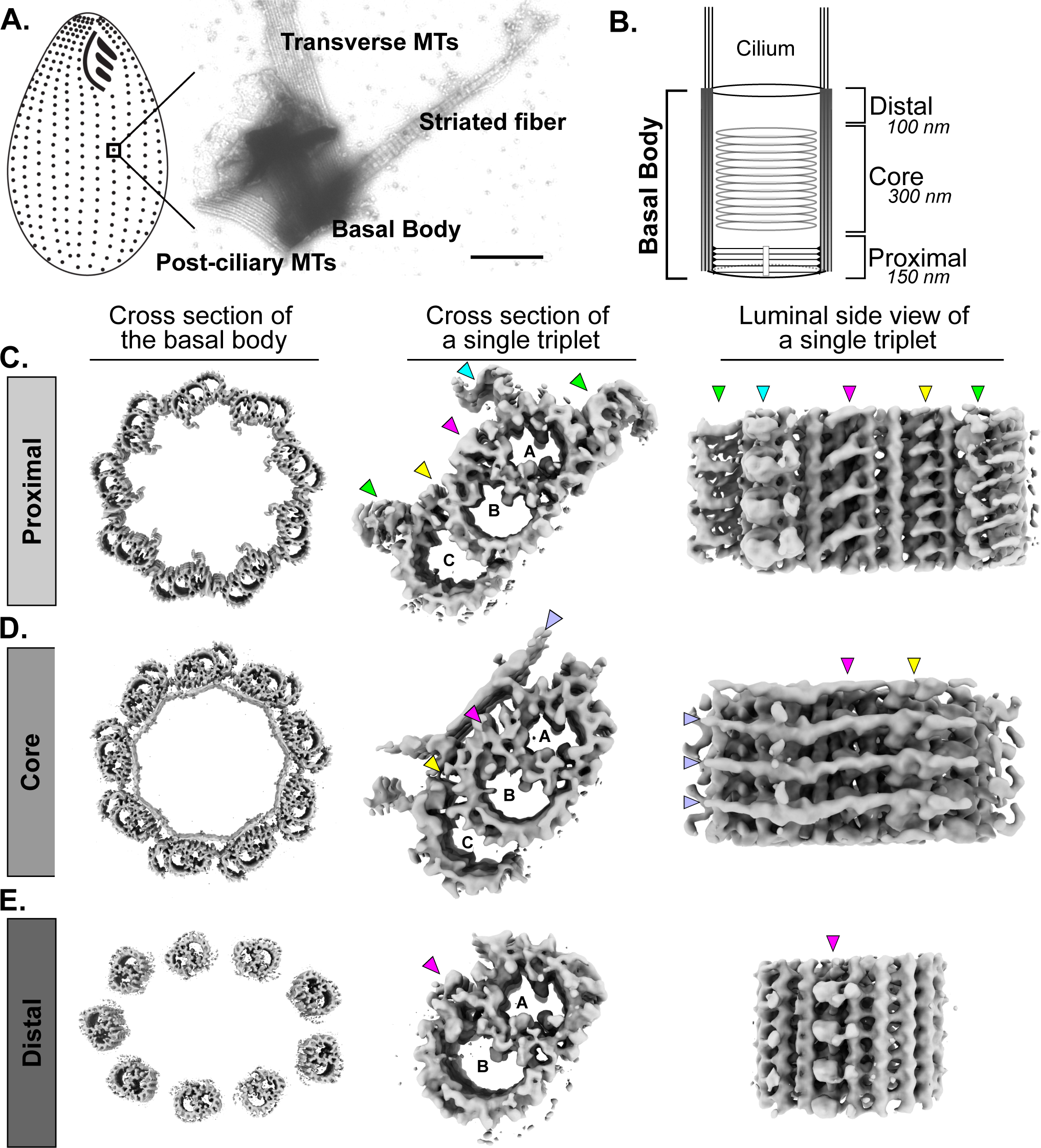
The structure of the *Tetrahymena* basal body. A) Cartoon diagram of a *Tetrahymena* cell with BBs in black on left. On right, negative stain image of a single BB isolated for cryoET showing BB appendage structures are maintained. Scale bar is 400 nm. B) Cartoon of the BB showing the three regions for which structures were determined: Proximal, Core, and Distal. The proximal region lumen harbors the cartwheel and the core contains the inner scaffold, indicated in the cartoon. C-E) CryoET structure of the *Tetrahymena* BB in the proximal (C), core (D), and distal (E) regions. Left: cross sectional view of a BB with the TMT subtomogram average mapped onto the TMTs in ice. Note that in the proximal and core regions the BB is circular, but the distal end is flattened. This is likely due to the force from freezing the BB on the EM grid in ice and a lack of an interconnecting structure at the distal end. Middle: cross sectional view of the TMT subtomogram average from each region. For the core region, this is the subclass of complete TMTs. For the distal region, no complete TMTs were observed. Right: luminal side view of the TMT subtomogram average structure. Arrow heads point to features of the TMTs: green, A/C linkers; cyan, pinhead; magenta, A-B inner junction; yellow, B-C inner junction; lavender, inner scaffold.

**Table 1.** Summary of Structures from Wild-type and poc1Δ Mutant.

| Structure | Number of Particles | Resolution (Å) | Description | EMDB ID # |
| --- | --- | --- | --- | --- |
| 1 | 7854 | 19.2 | Triplet MT in the proximal region, wild-type | EMD-42776 |
| 2 | 1467 | 27.7 | Triplet MT in the central core region, wild-type | EMD-42777 |
| 3 | 5670 | 19.2 | Doublet MT in the distal region, wild-type | EMD-42778 |
| 4 | 5834 | 9.82 | A/B inner junction in the proximal region, wild-type | EMD-42780 |
| 5 | 5071 | 10.0 | B/C inner junction in the proximal region, wild-type | EMD-42781 |
| 6 | 14373 | 8.47 | A/B inner junction in the central core region, wild-type | EMD-42782 |
| 7 | 868 | 25.3 | Triplet MT in the proximal region, poc1Δ | EMD-42783 |
| 8 | 1291 | 25.3 | Triplet MT in the central core region, poc1Δ | EMD-42784 |

By comparing our cryo-EM structures to those previously reported for other organisms, several consensus features can be identified. The proximal end structure of the TMT is highly conserved in *Tetrahymena*, *Chlamydomonas*, and *Paramecium* (Fig. S2A; (Li et al., 2019; Klena et al., 2020)). The positions and shape of the pinheads is similar among *Tetrahymena*, *Paramecium*, and *Chlamydomonas* BBs. In contrast, the A/C linkers are more variable in structure. The A/C linker in *Chlamydomonas* has an X-shaped configuration, but this is not the case in *Tetrahymena*, which lacks one leg of the X-shaped structure. Structural differences in the A/C linker are also observed between the *Paramecium* and *Tetrahymena* BBs, with the *Paramecium* A/C linkers appearing less dense. However, the degree to which these differences are species-specific or are due to the difference in resolution and / or flexibility is unclear (Fig. S2A). Side views of the TMTs from the BB lumen show high structural conservation between organisms. In particular, the structure and periodicity of the pinhead and densities of the A-B inner junction is similar; all exhibit 8 nm longitudinal periodicity, consistent with the periodicity of tubulin heterodimers in the microtubule lattice (Fig. S2A, bottom panels).

The core region and inner scaffold structure is more variable than the proximal region (Fig. S2B). *Tetrahymena* and *Paramecium* TMTs have primary attachment points to the inner scaffold at protofilament A03, and at the A-B and B-C inner junctions. However, the A03 attachment site is missing in the *Chlamydomonas* structure. In addition, the shape of the B-C inner junction attachment site is divergent compared to *Tetrahymena* and *Paramecium*. In *Tetrahymena* and *Paramecium*, the densities at the B-C inner junction are shorter and hold the scaffold more closely to the TMT inner walls, whereas in *Chlamydomonas* BBs, the B-C inner junction attachment densities create an elongated V-shape that holds the scaffold further away from the TMT walls. Despite these differences, the A-B inner junction attachment structure is nearly identical between all three species. We cannot rule out that the other attachments are simply more flexible in *Chlamydomonas* leading to their reduced density, or that they are invisible in this lower resolution structure. Regardless, the A-B inner junction is the most consistent attachment site in all three organisms compared here. In summary, the structure of the TMTs is broadly conserved throughout the BB, with most differences found in the core region. Within the core region, the attachment site of the inner scaffold at the A-B inner junction (the so-called “stem”; (Le Guennec et al., 2020)) is the most conserved attachment. The molecular composition of these structures and the mechanism of how they contribute to BB assembly and stability remain poorly understood.

### Poc1 is an inner junction protein in the *Tetrahymena* basal body

*Tetrahymena* poc1Δ cells assemble BBs and cilia that function under normal ciliary force conditions (growth at 30°C), but disassemble at high force (growth at 38°C; Fig. S3A; (Pearson et al., 2009a)). Inhibition of ciliary beating using NiCl_2_ rescues BB loss in *poc1Δ* cells at high temperature, verifying that the disassembly is due to ciliary forces (Fig. S3A; (Meehl et al., 2016)). To understand how the Poc1 stability protein binds in the BB architecture, Ultrastructure Expansion Microscopy imaged with Structured Illumination Microscopy (UExM-SIM) was used to determine Poc1 localization within BBs. In cross-sectional views, inducible GFP:Poc1 expressed in poc1Δ cells localizes to the interior TMT walls of BBs, creating a peak of ring-shaped intensity just interior to the peak of ring-shaped tubulin signal (Fig. 2A, top images). Longitudinal views of BBs revealed that GFP:Poc1 localizes through the length of the BB, to include the proximal and core regions, with a slight reduction of signal between the two domains (Fig. 2A, bottom images). Similar localization was observed using endogenously expressed C-terminal tags of Poc1, with the exception that clear rings were not observed in cross sectional views, suggesting either that the C-terminus is located interior to the N-terminus in the BB, or that it is dynamic, resulting in diffuse signal (Fig. S3B). Previous immunoEM studies showed localization of Poc1 through the length of the BB, consistent with the localization pattern observed here (Pearson et al., 2009a). In summary, *Tetrahymena* Poc1 localizes to the BB proximal and core regions, indicative of functions in both BB regions.

**Figure 2:**
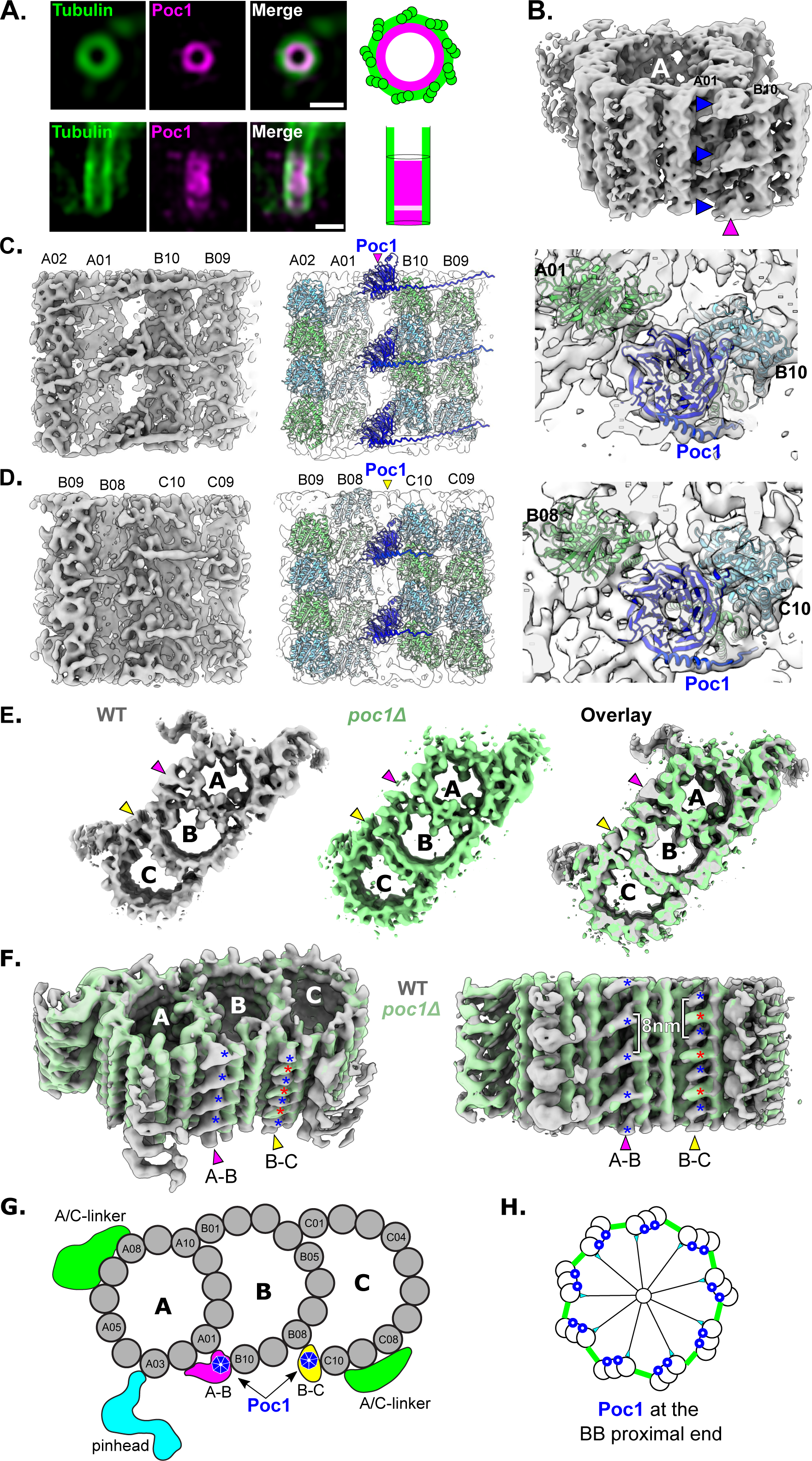
Poc1 localizes to the A-B and B-C inner junctions of triplet microtubules at the proximal end of the basal body. A) Cross sectional (top) and longitudinal (bottom) views of UExM-SIM expanded BBs from intact *Tetrahymena* cells, stained with anti-α-tubulin and anti-GFP antibodies to detect GFP:Poc1. The cells are *poc1Δ* rescued with GFP:Poc1. Scale bars are 500 nm, not corrected for expansion factor. B) Focused refinement of the WT A-tubule and A-B inner junction (magenta arrowhead) at ∼13 Å shows a doughnut shaped density (blue arrowheads) with 8 nm periodicity bridging protofilaments A01 and B10. C) The A-B inner junction density is a WD40 domain. Left: focused refinement of the A-B inner junction at 9.8 Å resolution, luminal side view. Middle and right show luminal side and top-down views, respectively, with ribbon diagrams of α- and β-tubulin (light green and cyan, PDB ID 8G2Z) modeled into protofilaments A01 and B10. An AlphaFold2 predicted structure of Poc1 is modeled into the A-B inner junction doughnut shaped density. See also Video 2 and Materials and methods for fitting. D) Focused refinement of the B-C inner junction at 10 Å, shows a WD40 density with 8 nm periodicity, interspersed with another protein density. Ribbon diagrams of α- and β-tubulin and Poc1 were modeled into the electron density as described above. E) Individual structures and overlay of WT TMTs (gray) with poc1Δ complete TMTs (green) in cross sectional view. Arrows indicate the positions of the A-B and B-C inner junctions and show a loss of density in poc1Δ TMTs. F) Glancing cross sectional view on left, luminal side view on right. The WD40 densities described at the WT A-B and B-C inner junctions (blue asterisks) in C and D are missing in the poc1Δ complete TMT, while the interspersed density at the B-C inner junction (red asterisk) remains. G) Schematic representation of the WT TMT structure from the proximal region of the *Tetrahymena* BB. Color scheme: gray, tubulin; green, A/C linker; cyan, pinhead; magenta, A-B inner junction; yellow, B-C inner junction; blue, Poc1. H) Schematic representation of Poc1’s localization at the proximal region A-B and B-C inner junctions of *Tetrahymena* BBs. Poc1 is indicated in blue.

To identify the precise localization of Poc1 in the cryoET structure, we searched for Poc1’s WD40 domain in the BB proximal region subtomogram average, focusing on the BB luminal side of the TMTs. We expected Poc1’s density to be shaped like a doughnut due to the seven-bladed β-propeller WD40 predicted structure. Indeed, a doughnut shaped density was identified at the A-B inner junction (Fig. 2B). Focused refinements on the A-B and B-C inner junctions substantially improved the resolution of the structure in these regions. Both inner junctions show a WD40 domain density where seven β-propeller blades can be resolved (Fig. 2C, 2D). Rigid body fitting of the AlphaFold predicted *Tetrahymena* Poc1 WD40 domain (UniProt ID Q229Z6) shows it fits well in the density (Fig. 2C, 2D, Video 2, see Methods).

To determine whether these WD40 densities were indeed the Poc1 protein, we prepared BBs from *Tetrahymena* poc1Δ cells using the BB isolation procedure described above (Video 3). The subtomogram average of poc1Δ BBs revealed a variety of TMT defects compared to the WT structure, including varying degrees of B- and C-tubule loss throughout the length of the BB. However, focusing on the subclass of complete TMTs from poc1Δ BBs showed that the WD40 densities at the A-B and B-C inner junctions were lost when compared to the WT structure, indicating they are indeed the Poc1 protein (Fig. 2E, F). In both inner junctions, Poc1 displays 8 nm longitudinal periodicity (Fig. 2F). In the A-B inner junction, Poc1 is the only protein found, whereas in the B-C inner junction, it is interspersed with an unknown protein that is retained in the absence of Poc1 (Fig. 2F). Thus, Poc1 localizes to the A-B and B-C inner junctions at the proximal end of *Tetrahymena* BBs (Fig. 2G, H).

Based on our highest resolution structures and by using the AlphaFold predicted model of Poc1 and previously determined α- and β-tubulin structure (PDB ID 8G2Z), we built a pseudo-atomic model for the A-B and B-C inner junctions, showing details of how Poc1 links the A- and B-tubules and the B- and C-tubules at the junctions (Fig. 2C, D, Video 2). In this model, the WD40 motif of Poc1 is tilted 45 degrees relative to the longitudinal direction of BB. The WD40 domain connects protofilaments A01 and B10 by arranging multiple copies of Poc1 protein every 8 nm longitudinally, effectively sealing the inner junctions of the TMT. Our model predicts that a loop in propeller blade two from Poc1’s WD40 domain binds to helix 12 of α-tubulin from protofilament A01. Meanwhile, blades three and four face protofilament B10, potentially making contact with an α-β-tubulin heterodimer in protofilament B10 at its longitudinal intradimer interface. In the A-B inner junction, a density that appears to be the C-terminal end of Poc1 exits the WD40 domain at blade number seven. It forms a long coil, horizontally traversing the luminal wall of the B-tubule, binding to protofilaments B09 and B08. A similar binding mode of the C-terminus exiting the WD40 domain is observed at the B-C inner junction. This long coil observed in the average density map is consistent with the predicted structure of the extended C-terminal tail of Poc1. In this model, the C-terminus of Poc1 at the A-B inner junction can be traced to approximately residue Q402 near protofilament B09. Residues 403-634 are not visible. The C-terminus of Poc1 at the B-C inner junction can be traced to residue Q375 near protofilament C10. A similar extended coil density that may be the Poc1 N-terminus exits the WD40 domain at the A-B inner junction, horizontally traverses a gap in the junction, and binds the luminal wall of the A-tubule at protofilament A02 (Fig. 2C, Video 2). Together, the subnanometer resolution structure refinement coupled with structural modeling reveals details of how Poc1 recognizes BB TMTs through the WD40 domain and potentially its N- and C-termini.

### Poc1 stabilizes triplet microtubules in the proximal region of the BB

Consistent with the *in vivo* BB instability found in poc1Δ cells grown in high-force conditions, isolated poc1Δ BBs displayed varying degrees of structural disintegration *in vitro* as assessed by negative stain EM, despite not having been exposed to high-force conditions, indicating their intrinsic structural instability (Fig. S3C, compare white arrowhead to red arrowhead). The cryoET and subtomogram averaging of poc1Δ BBs show partial B-tubules and missing C-tubules in the majority of BBs, inconsistent with prior EM-tomography studies showing poc1Δ BB TMTs *in situ* at normal-force are mostly complete (Meehl et al., 2016). This suggests that the tubule loss observed in the subtomogram averages likely occurred during BB isolation and freezing, as was observed for the WT BB in the core region. Subtomogram classification of poc1Δ TMTs in the proximal region showed substantial structural heterogeneity in different parts of the TMT. Only 8% of poc1Δ subtomograms contained complete TMTs with full A-, B-, and C-tubules. In contrast, 99% of WT BBs contained complete TMTs in the proximal region. This highlights that the poc1Δ structure is more labile than the WT structure, and differences in poc1Δ compared to WT BBs reveal the parts of the TMT that are sensitive to Poc1 loss, whether under forces from cilia or from BB isolation.

The direct binding of Poc1 to the TMTs suggested that Poc1 is required to promote closure of the B- and C-tubules against the A- and B-tubules, respectively, which in turn promotes TMT stability. Indeed, UExM-SIM of poc1Δ BBs in whole cells showed loss of entire TMTs (Fig. 3A). Consistent with these observations and as discussed above, only 8% of poc1Δ TMT subtomograms contained complete TMTs. Our cryo-EM tomography and subtomogram averaging therefore provides the opportunity to dissect this MT loss in more detail. Classification of the poc1Δ subtomograms shows various degrees of loss of the B- and C-tubules (Fig. 3C, D, Fig. S3E). No defects in the A-tubule protofilaments were observed. Protofilament loss from the B- and C-tubules occurs closest to the inner junctions. Classification focused on the B-tubule shows loss of the B10 protofilament that is immediately adjacent to the inner junction in most classes, as well as protofilaments B9 and down in protofilament numbering as the severity of the structural disintegration increases (Fig. 3C, classes B3-6). When all B-tubule protofilaments are present, two subclasses accounting for approximately half of the subtomograms were observed (Fig. 3C, classes B1 and B2). The major difference between these two subclasses resides in densities at the A-B inner junction both in the B-tubule lumen and the BB luminal side, where different proteins structurally distinct from Poc1 appear to keep the B-tubule attached to the A-tubule. In all classes, densities for the C-tubule are weak, indicating that C-tubule loss precedes B-tubule defects (Fig. 3C, all classes). Classification focused on the C-tubule reveals, as with the B-tubule, subclasses with protofilament loss from the inner junction and / or flexibility of those protofilaments, indicated by a smeared signal (Fig. 3D, classes C4 and C5). Three classes of complete TMTs were identified and combined to generate the poc1Δ complete TMT subtomogram average used in Fig 2 (Fig. 3D, classes C1-3). In summary, both UExM-SIM and focused classification of TMT subtomograms show that Poc1 in the proximal region promotes protofilament attachment at the inner junctions, thereby stabilizing TMTs directly.

**Figure 3:**
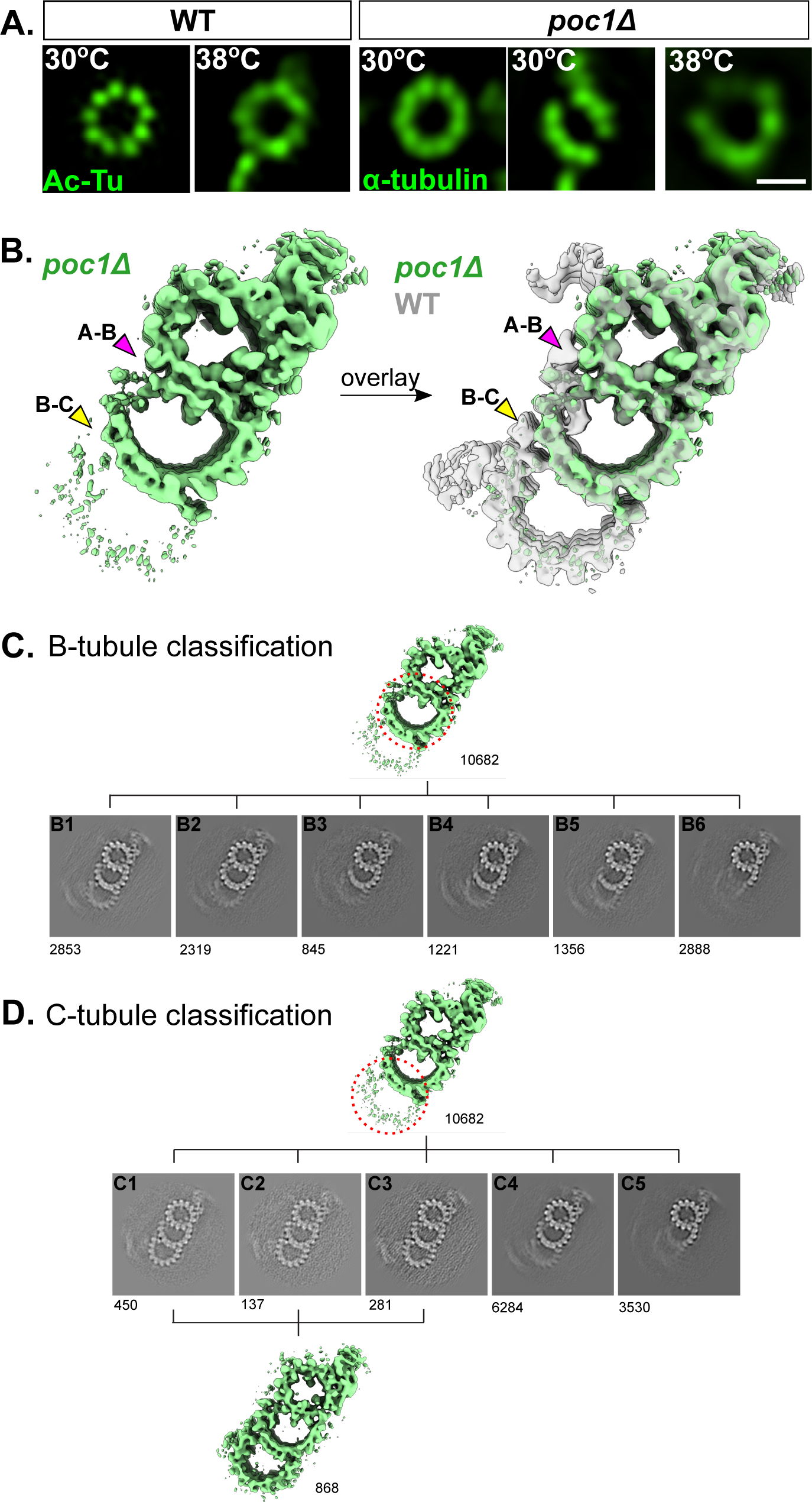
Poc1 stabilizes microtubules in the proximal region of the basal body. A) WT and poc1Δ BBs from whole cells expanded and imaged by UExM-SIM, stained with either anti-acetylated tubulin or anti-α-tubulin antibodies. Individual TMTs are observable in both WT and poc1Δ BBs. WT at 30°C and 38°C maintain all nine TMTs and overall circular shape. poc1Δ BBs are mostly comprised of nine TMTs at 30°C, but occasionally show loss of one or two TMTs (loss of two TMTs shown here). At 38°C, poc1Δ BBs lose uniform shape and tubulin signal is uneven, indicative of tubulin loss. Scale bar is 500 nm. B) Overall subtomogram average of poc1Δ TMTs in the proximal region shows loss of protofilaments. Left, poc1Δ TMT only (green). Right, poc1Δ overlaid with WT (gray). C) Classification of poc1Δ TMTs focused on the B-tubule (demarcated by a dashed red circle) into six classes displaying different states of B-tubule disintegration. The most severe protofilament loss is at the luminal side of the triplet, consistent with Poc1’s binding location on the TMT. The numbers of subtomogram in the classes are indicated . D) Classification of poc1Δ TMTs focused on the C-tubule (demarcated by a dashed red circle) into five classes displaying different states of C-tubule disintegration. Classes 1-3 are complete TMTs. The combined average of these classes was used for density loss comparison in Fig. 2E and Fig. 4B. The numbers of subtomogram in the classes are indicated.

### Poc1 stabilizes the pinhead in the BB proximal region

We next asked whether Poc1 stabilizes additional structures of the BB proximal end. A/C linkers were often disrupted in *poc1Δ* mutants even though detectable cartwheel formation and general BB assembly was unperturbed (Meehl et al., 2016). In both WT and poc1Δ complete TMT subtomogram averages, A/C linkers are present, but their density is smeared (Fig. 2E, F). This indicates flexibility or heterogeneity of the structure, even in WT BBs, making it difficult to assess quantitatively or qualitatively the differences between these structures in WT versus poc1Δ BBs. Thus, consistent with previous reports, A/C linkers can form in the absence of Poc1 protein {Meehl:2016gr}.

The pinhead links the cartwheel structure to the TMTs. A primary component of the pinhead is the α-helical protein, Bld10 / CEP135 (Hiraki et al., 2007; Lin et al., 2013; Guichard et al., 2017). To determine whether Poc1 impacts the localization of *Tetrahymena* Bld10, Bld10:mCherry levels were measured in both WT and poc1Δ cells (Fig. 4A). No difference was detected in the amount of Bld10 localized to WT versus poc1Δ BBs at normal cilia force conditions (30°C). However, under high-force conditions created by shifting cells to 38°C for 12 hours, Bld10 levels were reduced by approximately 60% at BBs in poc1Δ cells (Fig. 4A). Thus, Poc1 is not required for Bld10 assembly at the BB but does stabilize Bld10 in the face of elevated ciliary force.

**Figure 4:**
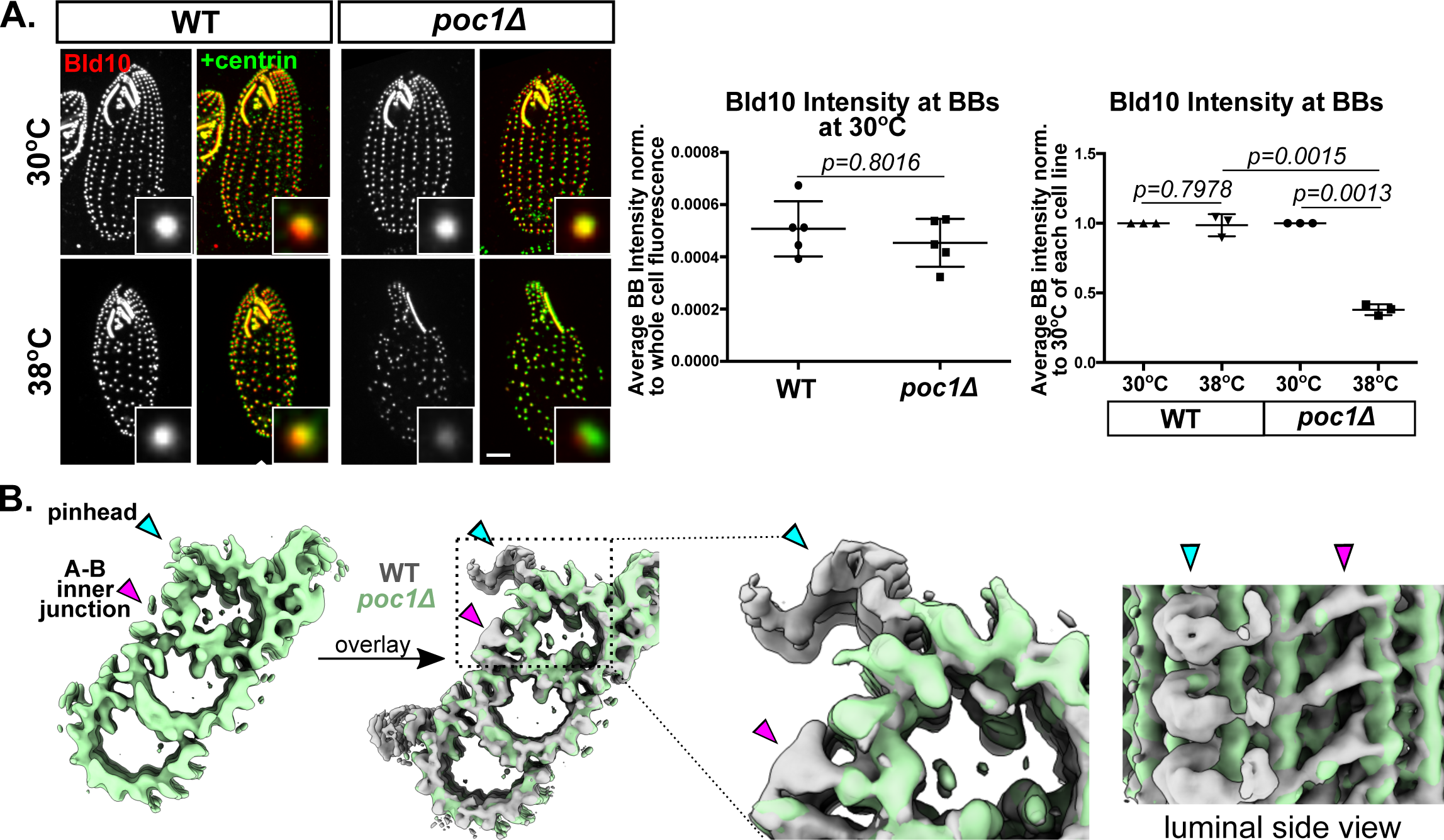
Poc1 stabilizes the pinhead at the proximal end of the BB. A) Bld10 is lost in high force conditions from poc1Δ BBs. Left images: WT and poc1Δ cells expressing Bld10:mCherry at 30°C and 38°C. Cells were fixed and stained with anti-mCherry antibody to visualize Bld10 (grayscale, red) and anti-centrin antibody to visualize all BBs (green). Cells were starved for 12 h to attenuate new BB assembly and temperature shifted for 12 h before fixation. Scale bar is 5 μm. Middle: Comparison of BB Bld10:mCherry levels at 30°C between WT and poc1Δ cells. Each point on the graph represents the average of 15 BBs, normalized to the total cell levels of mCherry determined for that cell (see Materials and Methods). There is no stastical difference in amounts of Bld10 at BBs between WT and poc1Δ cells. Graph shows mean +/− standard deviation. Right: Comparison of BB Bld10:mCherry levels in WT and poc1Δ cells at 30°C and 38°C. Bld10mCh levels at 30°C for each cell line were normalized to 1. Each point on the graph represents an independent experiment consisting of 150 BBs. Graph shows mean +/− standard deviation. BBs lose Bld10 at 38°C in poc1Δ cells, but not in WT cells. B) Cross sectional view of poc1Δ complete TMTs (green) alone and overlaid with WT TMTs (gray) shows loss of electron density at the pinhead. Right, zoomed in view of overlaid structure in cross-sectional and luminal side views. The pinhead is lost in poc1Δ TMTs even though the A-tubule structure where it should attach is intact. Cyan arrowhead, pinhead; magenta arrowhead, A-B inner junction.

The loss of Bld10 at elevated force in poc1Δ BBs led us to ask whether the structure of the pinhead was impacted by loss of Poc1. Indeed, the pinhead density was reduced in poc1Δ complete TMTs compared to WT TMTs (Fig. 4B). Thus, despite not being a primary component of the pinhead, Poc1 loss causes the pinhead to be unstable.

In summary, Poc1 stabilizes the proximal region of BBs. First, Poc1 stabilizes TMT tubule connectivity by binding directly to the MT protofilaments in the A-B and B-C inner junctions, sealing the tubules together. Second, Poc1 promotes stability of the pinhead structure that connects the TMTs to the cartwheel. This ultimately organizes and stabilizes TMT interconnections in the BB proximal region.

### Poc1 forms the base of the A-B inner junction attachment to the inner scaffold

Given our observation that Poc1 localizes in the BB core region by UExM-SIM, we studied its contribution to the BB core structure, function, and molecular composition. *Tetrahymena* homologs of the centriole and BB core region localizing proteins, Fam161A and Poc16/WDR90 were identified using BLAST analyses to the human proteins. Fam161A is an inner scaffold protein that binds to MTs and interacts with Poc1B in human centrioles and in yeast two-hybrid studies (Roosing et al., 2014; Le Guennec et al., 2020). Poc16 also localizes in the core region but is not thought to be an inner scaffold protein (Steib et al., 2020). Endogenous, mCherry tagged Fam161 and Poc16 localized to *Tetrahymena* BBs. Both proteins colocalized with Poc1 in the core region (Fig. 5A). In addition to its localization in the core region, Fam161A localized to the BB distal end, at what is assumed to be the transition zone at the cilium base (Fig. 5A, S5A). The distal localized population of Fam161A did not co-localize with Poc1. Thus, like Poc1, Fam161A and Poc16 localize to the core region in *Tetrahymena* BBs.

**Figure 5:**
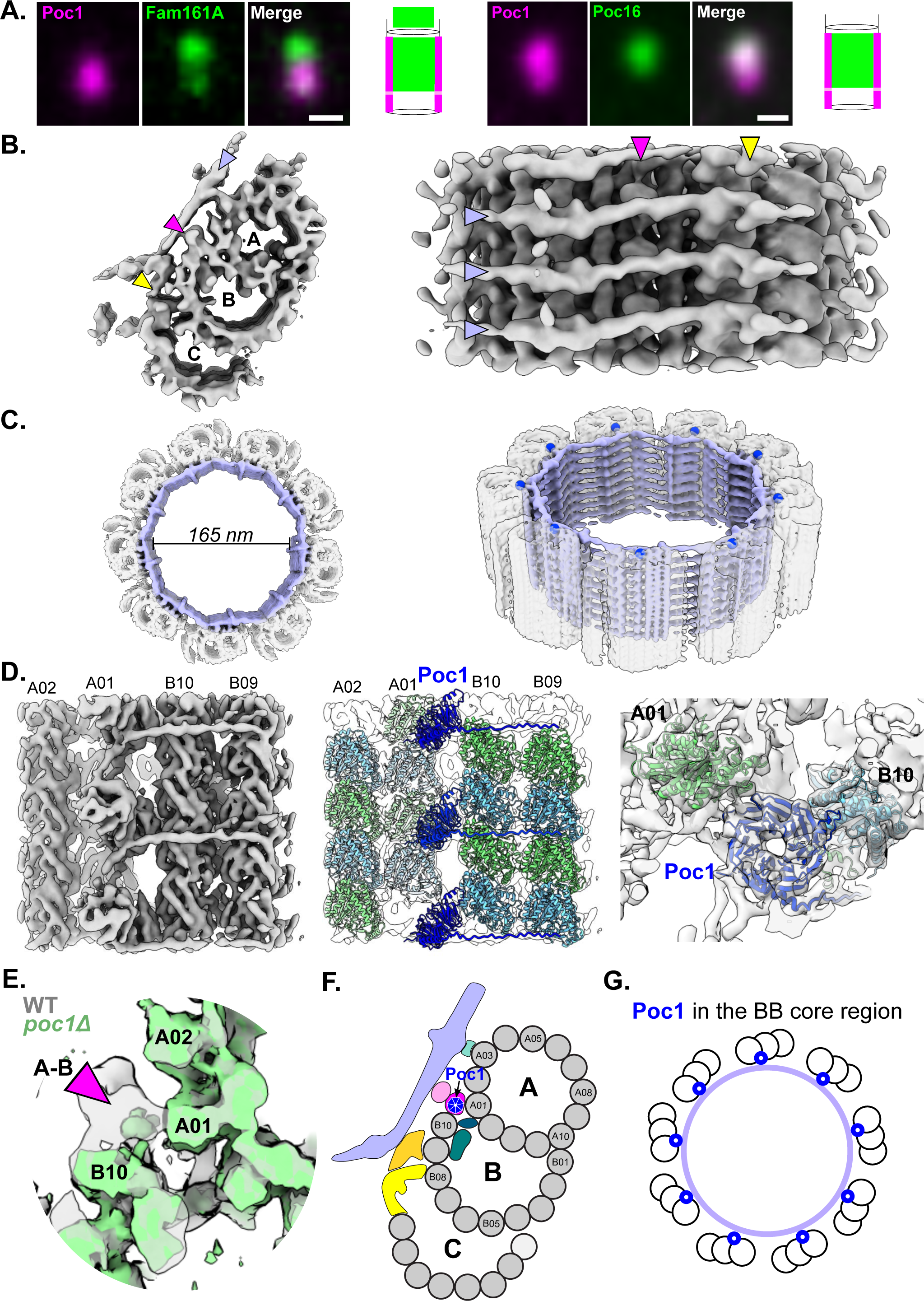
Poc1 is an anchor for the inner scaffold to the TMTs in the core region. A) Core region proteins Fam161A and Poc16 localize to BBs in *Tetrahymena*. Left, longitudinal view of Fam161A:mCherry (green) localization relative to Poc1:HaloTag (magenta) in WT cells. Right, longitudinal view of Poc16:mCherry (green) localization relative to Poc1:HaloTag (magenta) in WT cells. Longitudinal views are oriented with the cell exterior at the top of the image and the cell interior at the bottom. Diagrams of each protein’s localization are shown at right. Poc16 localizes to the core region only, whereas Fam161A has core and presumed transition zone localization. Images were acquired with a confocal microscope. Scale bars are 500 nm. B) Structure of the WT TMT from the core region. Left, subclass of complete TMTs (see Fig. S4A for overall average). Right, luminal side view of complete TMT. Arrowheads indicate the inner scaffold (lavender) and position of the inner scaffold attachment to the TMT at the A-B inner junction (magenta) and B-C inner junction (yellow). C) Complete TMT subtomogram average mapped onto an individual BB in ice shows the inner scaffold (lavender) and cylindrical shape. The A-B inner junction is indicated by blue dots in the image on right. The scaffold is a structure of stacked rings of 165 nm in diameter. D) Focused refinement on the A-B inner junction in the core region at 8.47 Å in luminal side view (left and middle) and top-down view (right). Ribbon diagrams for α- and β-tubulin and Poc1 are modeled into the density and colors are as described in Fig. 2 legend. An unknown protein density associates with the BB lumen-exposed side of Poc1 to form the A-B inner junction stem attachment to the inner scaffold. See also Video 4. E) Zoomed in view of the A-B inner junction from WT (gray) overlaid with poc1Δ (green) TMTs shows the WD40 density observed in the WT A-B inner junction (magenta arrowhead) is missing in the poc1Δ A-B inner junction. Both structures are from the subclasses of complete TMTs. F) Schematic representation of the WT TMT in the core region of the BB. Color scheme is as follows: gray, tubulin; lavender, inner scaffold; teal, A03 inner scaffold attachment; pink shades, A-B inner junction stem; yellow shades, B-C inner junction inner scaffold attachment; green shades, B-tubule luminal densities; blue, Poc1. G) Schematic representation of Poc1 (blue) localization in the BB core region with the inner scaffold represented by a lavender circle.

The core region is characterized by the inner scaffold, which is evident in our cryoET subtomogram average of *Tetrahymena* BBs (Fig. 5B and C). The overall structure of the inner scaffold is consistent with other organisms previously reported (Li et al., 2019; Klena et al., 2020; Le Guennec et al., 2020). In *Tetrahymena*, the inner scaffold repeat is an elongated structure attached to the luminal side of the TMT. It binds to the TMTs at multiple sites, as described in Fig. S2B; this includes protofilament A03 of the A-tubule, the A-B inner junction and the B-C inner junction. The inner scaffold exhibits 8 nm longitudinal periodicity. Laterally, it connects to the neighboring repeats near the A-tubule and the B-C inner junction of the TMT. Backfitting the core region TMT structure onto BBs revealed that, in *Tetrahymena*, the inner scaffold is comprised of stacked rings that circulate the luminal circumference of the BB (Fig. 5C, S4E). The density for the inner scaffold in WT BBs is not as well defined as the density for the TMTs themselves, indicating that the inner scaffold structure in our subtomogram average is heterogeneous or flexible. Nonetheless, our structure shows that the inner scaffold forms an interwoven meshwork in the lumen of the BB core region that holds the TMTs together in a manner that promotes structural integrity in the core region.

To determine whether Poc1 localizes to the A-B and B-C inner junctions in the core region, as found for the proximal region, we asked whether core region inner junctions also displayed WD40 densities. Focused refinement on the A-B inner junction revealed that, indeed, a density with seven β-propeller blades is present and an Alphafold model of *Tetrahymena* Poc1 can be docked into the structure (Fig. 5D, Video 4). The density was absent from the subclass of complete TMTs from poc1Δ BBs in the core region, validating that it is indeed the density for Poc1 (Fig. 5E). This suggests Poc1 localizes to the A-B inner junction in the BB core region as in the proximal region (Fig. 5E, 2C). Unlike in the proximal region, the BB core region B-C inner junction does not contain a Poc1 WD40 density. Rather, it is comprised of non-tubulin densities that are different from the Poc1-containing density found at the B-C inner junction at the BB proximal region (Fig. S4B-D). Moreover, comparison of the TMTs of poc1Δ BBs to WT TMTs at the B-C inner junction in the core region revealed no structural differences or lost density at the current resolution (Fig. S4D). This suggests that Poc1 is not a component of the B-C inner junction in the core of the BB. In summary, within the BB core region, Poc1 occupies the A-B inner junction, a major site of attachment of the inner scaffold to the TMTs (Fig. 5F,G).

### Poc1 stabilizes the BB inner scaffold

The A-B inner junction stem attachment is the most consistent structure linking the inner scaffold to TMTs across organisms and potentially serves as the major inner scaffold-TMT attachment site (Fig. S2B). Given Poc1’s localization at the A-B inner junction stem, we hypothesized that Poc1 influences the localization and stability of the Fam161A inner scaffold protein at BBs *in vivo*. Endogenously tagged Fam161A:mCherry localizes to BBs in poc1Δ cells under normal force conditions. However, this localization is decreased relative to WT cells (Fig. 6A, top panels). This is consistent with human Poc1B direct binding to Fam161A, but also shows that Fam161A loading in BBs is not entirely dependent on Poc1, suggesting there are redundant mechanisms to recruit Fam161A to BBs. When cells were shifted to high ciliary force conditions, Fam161A levels at the BB decreased by ∼60% relative to poc1Δ cells at normal force. A similar decrease in Fam161A levels was also observed in WT cells after temperature shift, suggesting that Fam161A is not stable in BBs at elevated ciliary force. Because Fam161A localizes to both the core and presumed transition zone BB regions (Fig. 5A), we asked whether one or both Fam161A populations are decreased in high ciliary force conditions. In both WT and poc1Δ cells, Fam161A was preferentially lost in the core region (Fig. S5A). This is consistent with Fam161A’s colocalization with Poc1 only in the core region.

**Figure 6:**
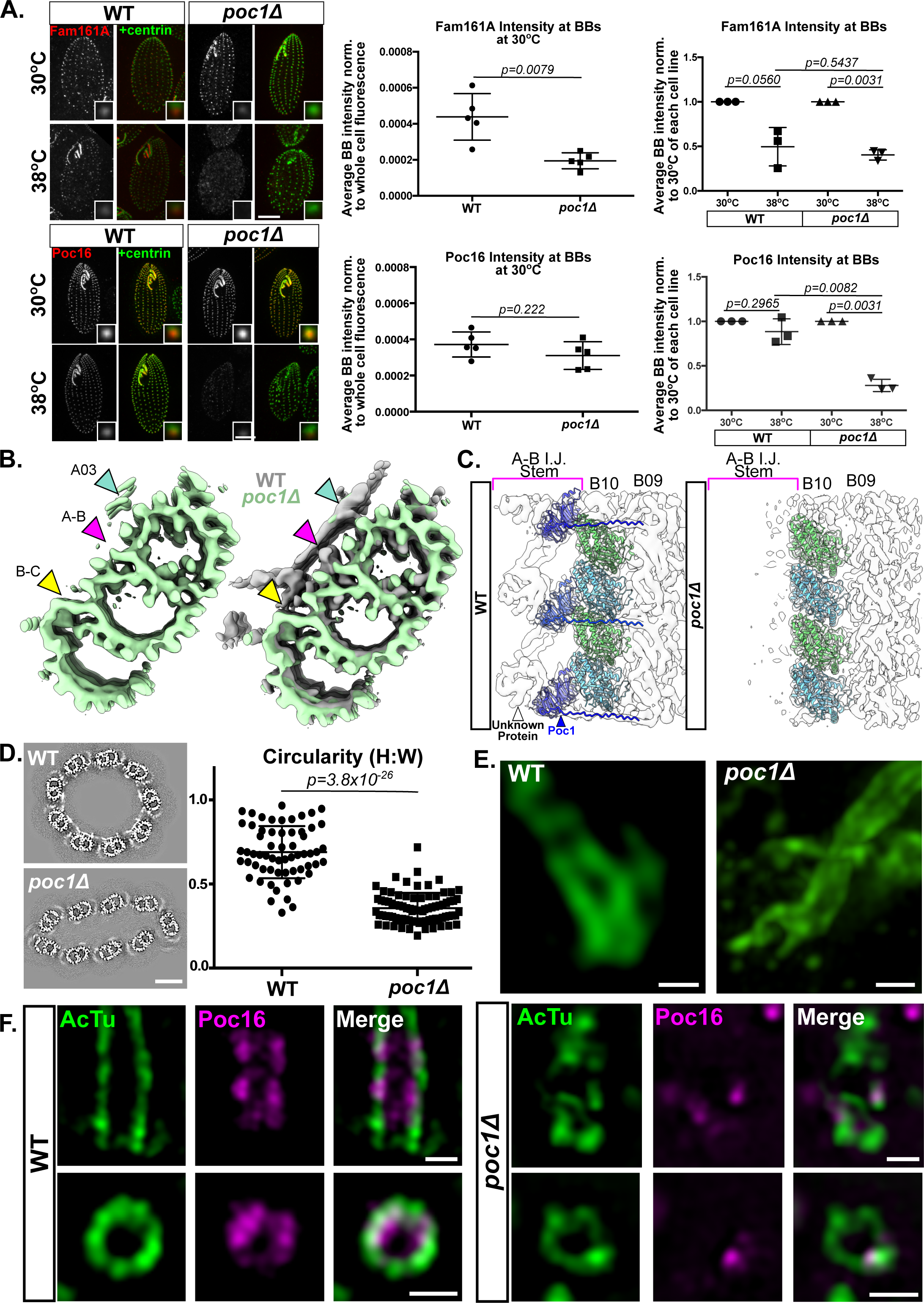
Poc1 stabilizes the inner scaffold in the core region of the BB and promotes BB shape maintenance. A) Fam161A and Poc16 localizations depend on Poc1 under high force conditions. WT and poc1Δ cells expressing Fam161A:mCherry (top left panels) and Poc16:mCherry (bottom left panels) at 30°C and 38°C. Cells were fixed and stained with anti-mCherry antibody to visualize Fam161A or Poc16 (grayscale, red) and anti-centrin antibody to visualize all BBs (green). Cells were starved for 12 h to attenuate new BB assembly and temperature shifted for 12 h before fixation. Scale bars are 10 μm. Middle graphs show quantification of Fam161A (top) and Poc16 (bottom) in WT versus poc1Δ BBs at 30°C, using total cell fluorescence to account for differences in assortment and / or expression between cell lines. BBs lacking Poc1 show defects in Fam161A, but not Poc16, binding under normal force conditions. Right graphs show quantification of Fam161A (top) and Poc16 (bottom) at BBs in WT versus poc1Δ cells at 30°C and 38°C. Values were normalized to the signal for each strain at 30°C. Each point on the graph represents an independent experiment consisting of 150 BBs. Graph shows mean +/− standard deviation. Both WT and poc1Δ BBs lose Fam161A upon shift to 38°C, whereas Poc16 is only lost in poc1Δ cells at 38°C. B) Cross sectional view of the complete TMT from the core region in *poc1Δ* alone (green) and overlaid with WT (gray). Arrowheads point to the inner scaffold attachment sites: A03 attachment site (teal), A-B inner junction attachment site (magenta), and B-C inner junction attachment site (yellow). Note the inner scaffold and A-B and B-C inner junction attachment sites are missing in poc1Δ TMTs, while the A03 attachment site is preserved albeit at a reduced level compared to WT. C) Side view of the focused refinement of the A-B inner junction / stem attachment in WT (left) and poc1Δ (right) TMTs showing in WT BBs Poc1 binds directly to the A-B inner junction and coordinates the binding of an unknown protein directly to Poc1’s WD40 domain. In poc1Δ BBs, both the Poc1 density and unidentified protein’s density are missing. D) Representative cross-sectional views of the core region of whole BBs in ice with the subtomogram averages from WT and poc1Δ TMTs superimposed on the TMTs from the individual BBs. Note that WT BBs are circular, whereas poc1Δ BBs are flattened. Bottom, graph of circularity measurements from all WT (n=61) and poc1Δ (n=81) BBs used in this study shows that poc1Δ BBs are significantly less circular than WT BBs. Graph shows mean +/− standard deviation. Scale bar is 50 nm. E) UExM-SIM of WT and poc1Δ BBs at 38°C stained with anti-α−tubulin antibody shows that WT BBs exhibit a gradual bend, whereas poc1Δ BBs display severe shape malformations that are likely to precede fracture and disassembly. Scale bar is 500 nm. F) UExM-SIM of WT and poc1Δ BBs at 38°C expressing Poc16:GFP and stained with anti-GFP (magenta) and anti-acetylated tubulin (green) antibody. Top panels show longitudinal views of individual BBs (with distal end of BB at top of the image) and bottom panels show cross sections of individual BBs. Poc16 is lost in poc1Δ BBs in places where tubulin defects are observable. Scale bars are 500 nm.

To determine whether Poc1 influences Poc16 localization to BBs, we visualized Poc16:mCherry in WT and poc1Δ cells. Poc16:mCherry intensity at BBs was unchanged between WT and poc1Δ cells in normal force conditions (Fig. 6A, bottom panels). However, Poc16 levels decreased by ∼70% in poc1Δ BBs at elevated ciliary force but were unchanged in WT BBs. These results suggest that Poc1 is partially required for Fam161A, but not Poc16, binding in the core region of the BB under normal force conditions, and that Poc1 stabilizes both Fam161A and Poc16 at the BB under high ciliary force conditions.

Because Fam161A was decreased in the core region in poc1Δ cells, we asked whether the inner scaffold is compromised in the poc1Δ BB structure. Indeed, a near complete loss of the inner scaffold was observed in the poc1Δ TMT structure from the core region, even when comparing the subclass of complete TMTs from both WT and poc1Δ BBs (Fig. 6B). The only remaining density appears at the A03 attachment site and is blurry compared to the MTs. This indicates decreased occupancy and/or increased flexibility of the molecules in that region. Therefore, in accordance with our fluorescence data, Poc1 stabilizes the inner scaffold.

To understand how the A-B inner junction attachment specifically was impacted by loss of Poc1, we characterized the structure of the stem at higher resolution. In WT BBs, the Poc1 WD40 motif density in the A-B inner junction directly binds another protein of unknown identity with a barrel shaped tertiary fold (Fig. 6C, left). This second protein attaches to the elongated horizontal density of the inner scaffold. In the absence of Poc1, the barrel density is missing, suggesting that Poc1 directly recruits and / or is necessary for the stable binding of this barrel shaped protein at the A-B inner junction to form the attachment site of the scaffold (Fig. 6C, right). We conclude that Poc1 stabilizes and organizes the inner scaffold by using its WD40 domain to create the base of the A-B inner junction attachment site, which then anchors the scaffold to the TMTs.

### Poc1 and the inner scaffold provide mechanical support to reinforce BBs

To understand how an intact inner scaffold contributes to the structural integrity of the BB, we asked whether there were gross differences in the shape of BBs from WT versus poc1Δ cells. Mapping the TMT subtomogram averages from WT and poc1Δ BBs onto individual BBs revealed that poc1Δ BBs, compared to WT BBs, were significantly flattened on the EM grid (Fig. 6D). The observed flattening was likely induced during BB isolation in centrifugation steps or by the compression force applied to the sample during blotting and freezing the EM grid, which was previously observed in centrioles from CHO cells (Greenan et al., 2018). This suggests that Poc1 and the inner scaffold bolster a sturdy architecture that is resistant to mechanical forces, including those introduced by cryoEM sample preparation.

We previously showed that BBs bend in response to the cilia beat stroke, suggesting that BBs balance rigidity with plasticity to ensure they do not disassemble (Junker et al., 2022). To understand how the BB architecture protects itself from ciliary forces, we used UExM-SIM to assess how BBs bend in response to ciliary beating. Under high force conditions, WT BBs exhibit a gradual bend, whereas dramatic fracturing of the BB through the core region was observed in the absence of Poc1, suggesting poc1Δ BBs lack the structural integrity that can integrate the bending without destructive deformation (Fig. 6E; (Junker et al., 2022)). To determine whether BBs that show structural defects have intact core structures, we used UExM-SIM to visualize Poc16 in BBs in high force conditions. In WT cells, Poc16 colocalizes with the tubulin walls of BBs as has been previously reported (Fig. 6F; (Steib et al., 2020)). However, in poc1Δ cells, Poc16 is sparsely found in the core region and is no longer closely associated with the TMTs. Regions lacking Poc16 display poor or absent TMT structure (Fig. 6F, S5B). We conclude that Poc1 contributes to BB structural integrity by promoting the assembly and / or stability of TMT interconnecting features in the BB core region.

## DISCUSSION

This study provides a structural mechanism to explain how Poc1 stabilizes BBs. Poc1, via its WD40 domain, seals the inner junctions of the A-B and B-C tubules at the BB proximal end, and the A-B tubules at the BB central core. By linking MT protofilaments at these sites, Poc1 stabilizes TMTs directly. Moreover, Poc1 promotes the stability of the pinhead and the inner scaffold, which interconnect TMTs and organize the BB architecture, thereby imparting structural integrity to the BB in the face of forces from beating cilia. Although conventional EM shows that poc1Δ cells assemble BBs with apparently normal TMTs under low-force conditions, the weak connections between TMTs, revealed in this study, render BBs unstable when ciliary forces are elevated. We establish a high-resolution structure-function relationship to explain the role of Poc1 in the mechanics of BB stability.

### Poc1 stabilizes proximal structures of BBs

#### Poc1 localizes to the A-B and B-C inner junctions

Poc1 was found to localize to the A-B and B-C inner junctions at the proximal end of BB TMTs using cryoET and subtomogram averaging (Figs. 1 and 2). The structures of WT A-B and B-C inner junctions at subnanometer resolution identified WD40 domains with seven β-propeller repeats, consistent with Poc1’s predicted structure (Fig. 2C, Video 3). Although we did not identify other doughnut shaped densities likely to be Poc1 based on our analysis of the WT and poc1Δ BB structures, we cannot exclude other, non-inner junction Poc1 localizations. Nevertheless, Poc1’s localization at the inner junctions rectifies several observations. In particular, it was previously proposed that Poc1 may be an A/C linker protein, but Poc1 is not exclusively localized to the BB proximal end. It also resides in the core region, which does not harbor proximal A/C linkers. In addition, the previously proposed doughnut structure for Poc1 in the *Chlamydomonas* A/C linker remains present in our *Tetrahymena* poc1Δ structure, suggesting that it is not a Poc1 protein density (Fig. S3D). Finally, the *POC1* gene has been lost from the *C. elegans* genome, a species whose BBs are comprised of singlet MTs rather than TMTs, and therefore have no inner junctions, obviating the need to maintain *POC1* in its genome (Keller et al., 2009). Together, Poc1’s specific localization to inner junctions along the length of the BB fits existing reports for how Poc1 localizes in the BB.

*Tetrahymena* Poc1 localization has differences when compared to its localization in human centrioles. In RPE-1 cells, Poc1B localizes along the BB length like in *Tetrahymena*, whereas in U2OS cells, Poc1B is only present in the central core region (Le Guennec et al., 2020; Pearson et al., 2009a). These species- and / or cell type-specific differences in Poc1 localization may reflect differences in BBs that template motile cilia versus primary cilia, or that exist in a non-ciliated state. The above studies focused on Poc1B, and it is possible that the human Poc1A ortholog preferentially occupies the proximal end of human centrioles and Poc1B resides at the central core region, in a separation of function between the two human orthologs. These distinct localization patterns could exist in a WT state, even though Poc1A and Poc1B can compensate for one another when knocked out individually (Venoux et al., 2013). Regardless, Poc1 promotes BB integrity through TMT stabilization and interconnection via its binding to the inner junctions, which is likely to be a unifying principle across phylogeny.

#### Poc1 stabilizes the proximal TMT B- and C-tubules

TMT loss is a primary defect in poc1Δ BBs (Meehl et al., 2016). In poc1Δ BBs, we observed missing or partially missing B- and C-tubules (Fig. 3). However, complete TMTs are formed in poc1Δ BBs and BB assembly rates are unaltered in poc1Δ cells, suggesting TMT formation itself is not a primary defect in poc1Δ cells (Pearson et al., 2009b; Meehl et al., 2016). Therefore, we interpret the difference in TMT structure observed here as a state of TMT loss via disassembly, rather than incomplete TMT assembly or maturation. However, interconnecting structures between TMTs may not be completely assembled in poc1Δ cells. The TMT structure loss is akin to the BB disassembly observed in poc1Δ cells upon high mechanical force from beating cilia. Here, the observed tubule loss in the tomograms likely was introduced by isolation and freezing required for cryo-ET. Multiple states of TMT disassembly were identified by classification of the TMTs from poc1Δ BBs (Fig. 3C,D). With the exception of the complete TMT subclass (8% of subtomograms), all other subclasses show significant structural defects at the A-B and B-C inner junctions. These defects manifest as the loss of protofilaments close to the inner junctions, to varying extents and changes to MT curvatures, even in less severe TMT disassembly states. One interpretation of these data is that these subclasses could represent a temporal view of TMT disassembly, wherein protofilament loss begins at the inner junction, and spreads towards the exterior side of the TMT in the subclasses with more severe defects. These data suggest that Poc1 serves to maintain a tight association of the B- and C-tubules with their neighboring tubules. In poc1Δ subclasses with partial B- or C-tubules, the tubules hinge at the outer junction and splay open, showing that the overall curvature of the B- and C-tubules in the WT BB is maintained in part by Poc1 and its role in sealing the inner junctions and stabilizing the TMT.

#### Poc1 stabilizes the pinhead

The proximal end of the BB contains A/C linkers and cartwheel structures that interconnect TMTs. Although defects in A/C linkers in poc1Δ BBs were previously reported, whether there are differences between WT and poc1Δ A/C linkers based on our cryoET structure is difficult to ascertain due to a limited amount of data and structural flexibility in the region. However, we found that poc1Δ BBs partially lost the cartwheel pinhead, and its molecular constituent, Bld10 (Fig. 4). This was an unexpected result given Poc1’s inner junction localization; we did not observe any WD40-like densities in the pinhead that would indicate an additional site of Poc1 localization there. This structure is reduced even in BBs with complete TMTs. One explanation for this loss is that the structural integrity of adjacent TMTs influences the pinhead. If there is an adjacent, structurally defective TMT, the pinhead attached to the complete TMT could be affected. However, complete TMTs in poc1Δ BBs tended to cluster together. This suggests that the complete TMTs are surrounded by other complete TMTs. Another explanation for pinhead reduction is that Poc1 influences its stability from a seemingly distant location in the inner junctions. It is possible that the N- and C-termini of Poc1 extend away from the inner junctions and, like outstretched arms, interact with the pinhead and A/C linkers to stabilize them. This is ostensibly inconsistent with our prior work showing that Poc1 lacking the C-terminal tail was sufficient to localize to BBs in poc1Δ cells and rescue BB loss at high force, unless other structures like the inner scaffold, which depends more directly on the WD40 domain, were recovered and are sufficient for the rescue (Pearson et al., 2009a). Regardless, this hypothesis of outstretched termini is strengthened by the high-resolution structure at the A-B inner junction in the proximal region, which reveals extended densities that protrude from the WD40 domain, consistent with a model in which one of Poc1’s termini links with the pinhead (Fig. 2C, Video 3). These densities reach laterally across the TMT toward the A-tubule on one side, where the pinhead resides, and toward the C-tubule and A/C linker attachments on the other side. These coil densities are absent in poc1Δ BBs, but our resolution of the poc1Δ structure is lower than that of WT, making it difficult to accurately compare the thin densities that only became distinguishable at higher resolution in WT BBs. Additional experiments are needed to determine if these horizontal densities are truly the Poc1 N- and C-termini and whether they interact with the pinhead and A/C linkers. If these densities are not extensions of Poc1, it suggests some other protein stretches across the Poc1 WD40 domain, laterally stabilizing the TMT, and potentially binds the pinhead and A/C linker. An additional model to explain pinhead loss in the absence of Poc1 is that forces exerted onto the TMTs cannot be distributed in a manner compatible with maintenance of the pinhead structure. Importantly, these models are not mutually exclusive. In summary, Poc1 supports the overall pinhead structure that links TMTs to the cartwheel, which appears critical for resisting forces from ciliary beating.

### Poc1 at the BB central core promotes inner scaffold stability

In the BB central core, Poc1 localizes to the A-B inner junction, creating the base of a conserved attachment site of the inner scaffold to the TMTs, referred to as the stem (Fig. 5D). The *Tetrahymena* TMT structure shows that the stem is also comprised of another, unidentified protein that binds directly onto Poc1’s WD40 domain (Fig. 6C). This unidentified Poc1-interacting protein in the BB core region reveals a solenoid protein fold (Fig. 5D, Video 4) and appears to connect the inner scaffold to the TMTs. Future experiments are needed to identify and characterize this unknown protein.

In both WT and poc1Δ BBs, we observed loss of C-tubules in the core region, with only 9.3% and 6.0% of the TMTs being complete, respectively (Fig. S4A). That C-tubules were lost in WT BBs indicates that they are generally less stable than A- and B-tubules. Incomplete TMTs were nonetheless capable of maintaining an inner scaffold, indicating that the C-tubules themselves are not essential for retaining the inner scaffold structure. This is perhaps due to the non-tubulin B-C inner junction structure in the core, which is still present in the absence of C-tubules, that serves as another attachment site for the inner scaffold (Fig. S4D). Complete TMTs from poc1Δ BBs did not retain any inner scaffold density at the B-C inner junction attachment site, and the inner scaffold density was greatly reduced at the A-tubule attachment site, even though Poc1 does not localize to those attachments. Thus, we conclude that the A-B inner junction stem is likely the most critical attachment site for organization of the inner scaffold.

The BB and centriole A-B inner junction in the core region was proposed to be occupied by Poc16 / WDR90 (Yanagisawa et al., 2014; Steib et al., 2020; Hamel et al., 2017). Poc16 is predicted to contain a DUF667 motif at its N-terminus and two to four WD40 domains in the remainder of the protein, depending on the organism (Steib et al., 2020). This is similar to the structure of the cilia axoneme A-B inner junction protein Fap20, which is a DUF667 fold-containing protein, and the two WD40 domain-containing axonemal protein, Fap52, which resides in the B-tubule lumen. Thus, Poc16 was proposed to be a fusion of a Fap20-like domain with a Fap52-like domain that uses the Fap20-like domain to bind the inner junction (Yanagisawa et al., 2014; Steib et al., 2020; Hamel et al., 2017). However, the shape of the density at the *Tetrahymena* A-B inner junction is that of a 7-bladed WD40 domain and not a DUF667 domain (Fig. 5D, Video 4). In addition, an Alphafold prediction of *Tetrahymena* Poc16 shows a β-sandwich domain followed by four seven-bladed WD40 domains. This predicted structure does not fit into the densities at the A-B inner junction. Finally, Poc16 localizes to poc1Δ BBs *in vivo* while the cryoET structure shows the A-B inner junction doughnut and barrel structures to be completely absent (Fig. 6C). This suggests that, in *Tetrahymena*, Poc16 resides elsewhere in the TMT core architecture and that Poc1, but not Poc16, resides at the A-B inner junction of the BB core region. Further studies are needed to define Poc16’s localization in TMTs.

Fam161A is reported to be an inner scaffold protein and to bind microtubules and Poc1B (Le Guennec et al., 2020; Roosing et al., 2014). It is predicted to have long, unstructured regions with stretches of α-helices dispersed throughout. Exactly how Fam161A contributes to the inner scaffold structure is unknown. Our fluorescence data show that Fam161A localizes to BBs in the absence of Poc1, albeit at a reduced level (∼50% reduction). This suggests that Fam161A has redundant mechanisms to localize to the *Tetrahymena* BB core region. While the vast majority of the density for the inner scaffold was lost in poc1Δ BBs, some density remained at the protofilament A03 attachment site (Fig. 6B). We propose that in the absence of the A-B inner junction attachment established by Poc1, inner scaffold proteins attached to the A03 site are no longer organized horizontally across the TMT. This causes them to both be unstable and unlikely to reach binding partners near the TMT C-tubule. Whether Fam161A binds to the A03 attachment site or elsewhere is unknown, but in either case, the cryoET structure corroborates the fluorescence data showing reduction of Fam161A in the core region.

Two Fam161A populations are present in *Tetrahymena* BBs: one localizes at the core region and a second at the distal end, which we assume to be the transition zone (Fig. 5A, S5A). This distribution of Fam161A localization may be analogous to that found at the BB and connecting cilium of the human retina (Mercey et al., 2022). Fam161A intensity decreased in poc1Δ BBs and, surprisingly, in WT BBs when cells were shifted to high-force conditions (Fig. 6A, S5A). This may reflect a dynamic molecular and structural remodeling of the inner scaffold to improve force resistance, akin to force-responsive maturation and remodeling of focal adhesions (Grandy et al., 2023; Legerstee and Houtsmuller, 2021). Precedence for such remodeling is evident with Poc5, a proposed inner scaffold protein in humans, but which is only present in immature BBs in *Tetrahymena* (Heydeck et al., 2020). Further experimentation is needed to understand these putative remodeling events. Moreover, a greater understanding of the repertoire of inner scaffold proteins in *Tetrahymena* and other organisms is necessary to understand how they are used to create functional inner scaffolds.

Our work focused on BBs that template motile cilia and resist their mechanical forces during ciliary beating, but loss of human Poc1 disrupts centriole integrity outside of the context of motile cilia. For instance, Poc1B mutations have been linked to polycystic kidneys and cone-rod dystrophy, and Poc1A mutations cause dwarfism among other developmental defects (Shaheen et al., 2012; Sarig et al., 2012; Shalev et al., 2012; Roosing et al., 2014; Durlu et al., 2014; Beck et al., 2014). These clinical presentations implicate Poc1A and Poc1B in the function of BBs that nucleate primary cilia or centrioles that are not ciliated. In kidneys, fluid flow is sensed by the deformation of primary cilia, stimulating a mechanosensory-based signal transduction pathway (Praetorius and Spring, 2001). The deformation of the cilium likely imposes forces on the BB that must be stabilized to resist disassembly, defining a clear need for Poc1 in those BBs. Similarly, photoreceptor cells are mechanosensitive, and cone-rod dystrophy can result from a defective connecting cilium structure, which is defined in part by the connecting cilium inner scaffold (Bocchero et al., 2020; Mercey et al., 2022). Thus, the assembly of the connecting cilium inner scaffold may depend on the proper assembly of the Poc1-dependent centriolar inner scaffold. Finally, forces from the mitotic spindle and microtubule-dependent centrosome movement during G2/M are resisted, in part, by posttranslational modifications of centriolar TMTs (Abal et al., 2005; Bobinnec et al., 1998). Poc1 may further stabilize centrioles against these microtubule-dependent forces. Together, these observations suggest that Poc1 broadly functions to stabilize BBs and centrioles against forces even outside of the context of ciliary beating.

We show that ciliary force resistance imparted by Poc1 in the *Tetrahymena* BB is multifaceted. First, Poc1 binds the inner junctions of BB TMTs, linking protofilaments at the junctions and promoting tubule architecture that supports BB stability by preventing TMT disassembly. Second, Poc1 stabilizes interconnecting structural features and their attachments to the TMTs. This is exemplified by Poc1’s direct interaction with the inner scaffold via its WD40 domain in the core region, but also occurs through unclear mechanisms to stabilize the pinhead in the BB proximal region. Such functions of Poc1 lend insight to its strong conservation across phylogeny and devastating consequences when mutated in humans.

## MATERIALS AND METHODS

### *Tetrahymena* cell culture and growth media

*Tetrahymena thermophila* strains B2086 (TSC_SD01625) and SB1969 (TSC_SD00701) were obtained from the *Tetrahymena* Stock Center at Cornell University. Creation of the poc1Δ and Poc1:mCherry and GFP:Poc1 strains is described in (Pearson et al., 2009a). Cells were grown to mid-log phase at 30°C in 2% SPP (2% proteose peptone, 0.2% glucose, 0.1% yeast extract, and 0.003% Fe-EDTA) unless otherwise indicated. Starvation media was 10mM Tris-HCl pH 7.4. Live and fixed cells were collected by centrifugation at 0.5xg for 3 min unless otherwise indicated. Cell densities were determined using a Coulter Z1 cell counter with size gating of 15 to 45 μm.

### BLAST analysis

NCBI protein Basic Local Alignment Search tool was used to identify *Tetrahymena* homologs of human FAM161A (accession number NP_001188472.1) and WDR90 / Poc16 (accession number AAI21187.1). For FAM161A, we identified TTHERM_00052590, with an E-value of 5×10^−11^ and 29% sequence identity. For WDR90 / Poc16, we identified TTHERM_00218620, Hs>Tt 5×10^−55^ with 28.2% sequence identity.

### Plasmid construction

The following constructs were generated to create C-terminal appended fluorescent proteins for expression in *Tetrahymena* cells: Poc1:HaloTag (Neo2); Fam161A:mCherry (Blasticidin); Poc16:mCherry (Blasticidin); Poc16:GFP (*NEO2*). For Fam161A and Poc16 constructs, homology arms approximately 400 bp each, separated by a NotI restriction site, were synthesized in pUC57 at the EcoRV site by Genscript (Piscataway, NJ). The upstream homology arm sat at the end of the coding portion of the gene but eliminated the termination codon. Inserts containing mCherry or GFP and blasticidin or *NEO2* resistance genes were amplified by PCR from plasmid p4T2-1 (Winey et al., 2012) with the following primers: NotIp4T21US 5’-GCGCGGCCGCTAAAGAAACTGCTGCTGCTAAATTCG-3’ or NotI p4T2 GFP US F 5’-ACAGCGGCCGCtttaATGAGTAAAGGAGAAGAACTTTTCAC-3’ and NotIp4T21DS 5’-gcGCGGCCGCCTAACATGTATGTGAAGAGG-3’. The inserts and synthesized homology arm vectors were then digested with NotI, gel purified, and ligated using 2x Ligation Mix (Takara) and checked for insert orientation. The N-terminal GFP tag of Poc1 used in this study was created previously (Pearson et al., 2009a) and the Poc1:Halo tag construct was created by subcloning the HaloTag from p4T2-1 Sas4:HaloTag vector into the p4T2-1 Poc1:mCherry vector using EcoRI and XbaI cut sites (Ruehle et al., 2020; Pearson et al., 2009a). All plasmid sequences were verified using whole-plasmid nanopore sequencing technology (Plasmidisaurus; Eugene, OR).

### *Tetrahymena* strain production

Macronuclear transformation was performed as previously described using DNA-coated particle bombardment (Bruns and Cassidy-Hanley, 2000). Transformed clones were selected using 100 μg/mL paromomycin to select for the *NEO2* gene, 60 μg/mL blasticidin S for the *BSR* gene, or 7.5 μg/mL cycloheximide to select for the *CHX* gene. To increase the copy number of the endogenously tagged constructs, cells were assorted using increasing concentrations of the appropriate drug.

Cell lines created in this work expressing fluorescent fusion proteins include: Poc1:HaloTag in SB1969 background; Fam161A:mCherry in SB1969, SB1969 with Poc1:HaloTag, and *poc1Δ* □□□□□□□□□□□; Poc16:mCherry in SB1969 background, SB1969 with Poc1:HaloTag, and *poc1Δ* □□□□□□□□□□□; Poc16:GFP in SB1969 and *poc1Δ* backgrounds.

### Basal body preparation for cryo-tomography

*Tetrahymena* cells were grown in 200 mL SPP prepared with HPLC grade water to high-log cell concentrations (0.5 – 1.0 x10^6^ cells/mL) at 30°C. All media and buffers were made using HPLC grade water, as this was determined to be important for cell lysis. Cells were pelleted, washed with 1x PBS, then resuspended in cold 18.75 mL Osmo Buffer (10mM Tris pH 7.5, 1M sucrose, 1mM EDTA) containing freshly prepared 1mM phenyl methyl sulfonyl fluoride (PMSF) protease inhibitor and incubated on ice for 5 minutes. Samples were transferred to 50 mL conical vials and TritonX-100 was added to a final concentration of 7.5% plus 1 μl of neat β-mercaptoethanol. 7.5mL of sterile glass beads were added to the samples and the conical vials were vortexed 3 times with 4-second pulses. Vials still containing the glass beads were then nutated at 4°C for 1-4 hours, monitoring progress of the lysis with a brightfield microscope approximately every 30 minutes. Lysis was considered complete when whole cell bodies were largely disrupted and cytoplasmic streaming was observed. This process was complete at about 4 hours for wildtype cells and within about 1.5 hours for the *poc1Δ* cells. Upon completion of cell lysis, samples were diluted 3-fold in 1x TE buffer and spun in a JA25.2 fixed angle rotor at 13,000xg for 45 minutes to pellet the basal bodies. The pellet contained a looser, “fluffy” layer which, in addition to basal bodies, contained more cell debris and whole oral apparatuses, as well as a glassier, “hard” pellet, which contained more pure basal bodies, though other macromolecular complexes are found in this pellet too, including proteasomes and ribosomes, as determined by negative staining the samples. After carefully removing the supernatant and fluffy pellet, the hard pellet was washed with 500 μl of 1x TE buffer to remove any residual cell debris, and then was resuspended in 500 μl of 1x TE buffer. Cryo-EM grid preparation was performed immediately afterwards to prevent sample degradation. Freeze-thaw destroys the structural integrity of the sample.

### EM Grid preparation

Grids for cryoEM were prepared using 200 mesh Quantifoil Cu EM grids with 2 μm holes (Ted Pella). Grids were glow discharged for 1 minute in negative ion mode. 4 μl of basal body sample containing BSA-coated 10 nm gold fiducials (BBI Solutions) was applied to the grid and plunge frozen in liquid ethane after a 45 s wait time using a ThermoFisher Vitrobot. Relative humidity of the chamber was 95%, temperature was 22°C, and blot time was 0.5 s. Negative stains to assess sample quality were prepared immediately after freezing using 3 μl of BB sample on 400 mesh copper grids and staining with 3% uranyl acetate. Negative stained samples were imaged on a 120 kV ThermoFisher/FEI Talos L120C microscope with a Ceta CMOS detector.

### Cryo-electron tomography data collection

Single-axis tilt series were collected on two field emission gun 300 kV Titan Krios electron microscopes (Thermo Fisher, Inc) at UCSF. Each scope was equipped with a Bio-Quantum GIF energy filter and a post-GIF Gatan K2 or K3 Summit Direct Electron Detectors (Gatan, Inc.). The GIF slit width was set at 20 eV. SerialEM was used for tomography tilt series data collection (Mastronarde, 2005). The data were collected in the super-resolution and dose-fractionation mode. The nominal magnification was set at 33,000. The effective physical pixel size on recorded images was either 2.70 Å (for wildtype) or 2.65 Å (for *poc1* KO mutant). A dose rate of 20 electron/pixel/second was used during exposure. The accumulated dose for each tilt series was limited to 80 electron/Å^2^ on the sample. A bi-directional scheme was used for collecting tilt series, starting from zero degree, first tilted towards −60°, followed by a second half from +2° to +60°, in 2° increment per tilt.

### CryoET data processing and model building

For tomogram reconstruction and subtomogram averaging, the dose-fractionated movie at each tilt in the tilt series was corrected of motion and summed using MotionCor2 (Zheng et al., 2017). The tilt series were aligned based on the gold beads as fiducials by using IMOD and TomoAlign (Kremer et al., 1996; Fernandez et al., 2018). The contrast transfer function for each tilt series was determined and corrected by TomoCTF (Fernández et al., 2006). The tomograms were reconstructed by TomoRec taking into account of the beam-induced sample motion during data collection (Fernandez et al., 2019). A total of 63 wildtype BB tomograms from 61 tilt series, 85 *poc1Δ* BB tomograms from 83 tilt series were used for reconstruction and subtomogram averaging.

For subtomogram averaging, first the BBs were identified in the 6xbinned tomograms. The center of TMT and their approximate orientation relative to the tilt axis were manually annotated in a Spider metadata file (Frank et al., 1996). The initial subtomogram alignment and average were carried out in a 2x binned format (pixel size 5.4 Å for the wildtype or 5.30 Å for *poc1Δ* mutant). The longitudinal segment length of basal body TMTs in a subtomogram was limited to 24 nm and 50% overlapping with neighboring segments. Without using any external reference, the subtomogram alignment was carried out by a program MLTOMO implemented in the Xmipp software package (Scheres et al., 2009).

Since TMTs from BBs are a continuous filament, after obtaining the initial alignment parameters, a homemade program RANSAC was used to detect and remove any alignment outliers and to impose the continuity constraint on the neighboring segments. This corrected the misaligned subtomograms by regression. MLTOMO and Relion 3.1 or 4.0 were extensively used for focused classification of the subtomograms (Bharat and Scheres, 2016; Zivanov et al., 2022). This was critical for determining the correct periodicity of the MIPs and for identifying structural defects or heterogeneity in the TMTs. These out-of-register subtomograms were re-centered and re-extracted. This was followed by combining all subtomograms for the next round of refinement. The final refinements, focusing on the inner junctions of the TMT, were carried out by using a workflow implemented in Relion 4.0 (Zivanov et al., 2022). The overall resolutions were reported (Fig. S1, Table 1) based on the Fourier Shell Correlation (FSC) cutoff at 0.143 (Scheres and Chen, 2012; Rosenthal and Henderson, 2003).

The pseudo-atomic models for the inner junctions at different regions of the basal body were built in ChimeraX (Pettersen et al., 2021) by fitting previously published atomic models or models predicted by AlphaFold2 (Jumper et al., 2021) into the subtomogram averaging density maps. UCSF ChimeraX was also used for visualization and for recording images.

The final average maps, including their EMDB access codes are summarized in Table 1.

### Structural analysis on the Inner scaffold

Since the MT is a dominant feature in the averaged TMT structure, the overlapping MT signal could interfere with the analysis of the inner scaffold. To eliminate this potential problem, a soft-edged binary 3D mask was initially generated and imposed onto the TMT average. This removed MT backbone density from the average and kept only the inner scaffold structure. Based on the TMT subtomogram location and their 3D refinement parameters defined during the refinement, the masked average volume containing only the inner scaffold structure was then backfit into the tomograms. The backfitting took into account the translations needed to put the TMT into the register of its 16 nm periodicity in the inner junction region. The resulting model was a barrel-shaped structure containing only the inner scaffold.

To analyze the arrangement of inner scaffold, the inner scaffold volume calculated above was projected on a plane orthogonal to the longitudinal axis of the BB. This was followed by calculating the Fourier transform of the projection. The resulting Fourier showed characteristic layer lines, indicating the nature of its helical assembly. The Fourier had strong signal at 8 nm layer line, indicating the 8 nm periodicity of the inner scaffold (Figure S4F). A line scan along the 8 nm layer line showed the maximal intensity near the center, crossing the meridian, strongly indicating its Bessel order close to zero (DeRosier and Moore, 1970; Klug et al., 1958). In real space, this translates into the inner scaffold repeat unit forming a “zero start” helix, a closed circular ring. The inner scaffold is arranged as a stack of multiple rings that are 8 nm apart perpendicular to the BB longitudinal axis. An example process is illustrated in Fig. S4E. The analysis was repeated on 63 WT BB datasets at different longitudinal location in the core region, and had a consistent result.

To confirm the above conclusion, auto-correlations are calculated while rotating the BB longitudinal axis in 40-degree step on the regenerated inner scaffold volume that has nearly intact 9-fold symmetry. The nine correlation maxima have minimal translation along the BB longitudinal axis, indicating the repeat units have a rise close to zero. They are on the same plane perpendicular to the longitudinal axis.

Finally, manual tracing and inspecting connection is conducted by following the inner scaffold ring around the inner circumference of BB. It shows consistent results that the inner scaffold arranges as longitudinal stack of closed rings in *Tetrahymena* (Fig. 5C).

### BB protein level analysis in WT and *poc1Δ* cells in normal and high force conditions

Cells grown to mid-log density in 2% SPP were pelleted and resusupended in 10mM Tris-HCl pH 7.4 starvation medium and distributed equally between two flasks for each cell line. Both flasks for each cell line were incubated at 30°C for 12 h, then one flask of each cell line was temperature shifted to 38°C. After 12 h, the cells both at 30°C and 38°C were harvested and stained by immunofluorescence as described below. For experiments with NiCl_2_ treatment, starved cells were treated with 300 μM freshly prepared NiCl_2_ for 1 h at 30°C to verify inhibition of ciliary beating before shifting to 37°C for 16 h. Image analysis was performed as follows: maximum projected images through half a cell volume were generated for each image. Fifteen BBs from 10 cells (150 BBs total) were analyzed for each condition in each replicate. To determine the signal from each BB, a 0.65 x 0.65 μm square region of interest was centered over the BB signal of interest, followed by two squares of the same size placed immediately next to the BB to determine local background signal. Integrated densities of all boxes were measured and the average of the two background boxes was subtracted from its corresponding BB box to determine the fluorescence value of that BB. The average value of BB fluorescence at 30°C was normalized to 1 for each cell line. Three biological replicates were performed. To determine the amount of protein at BBs in WT versus poc1Δ cells, sum-projected images of the entire cell volume were created and a custom region of interest was drawn tightly around the cell edge. Five 25 x 25 μm squares were placed around each cell to measure local background signal. Integrated densities and areas of each region of interest were measured and integrated density per area values were calculated. The average integrated density per area of the background boxes was subtracted from the cell’s integrated density per area, then multiplied by the cell’s area to calculate the total cell fluorescence signal. These values were then used to normalize the BB fluorescence intensities determined above for 15 BBs in five cells per cell line at 30°C. This step was necessary to account for slight differences in assortment level between cell lines.

### Immunofluorescence

Cells were pelleted and fixed in 3.2% paraformaldehyde 0.5% TritonX-100 in 1x PHEM buffer for 10 min at room temperature. Fixed cells were then pelleted, washed twice with 0.1% BSA-PBS and incubated in primary antibody solution in 1% BSA-PBS either for 2 h at room temperature or overnight at 4°C. Cells were then washed 3 times with 0.1% BSA-PBS and incubated in secondary antibody solution in 1% BSA-PBS either for 1 h at room temperature or overnight at 4°C. Cells were washed twice in 0.1% BSA-PBS then once in 1xPBS before mounting on coverslips with Citifluor AF1 mounting medium (Citifluor; Hatfield, PA). Coverslips were sealed to the slides using clear nail polish. Primary antibodies used in this work are documented in Table 2. Secondary antibodies were derived from goat and conjugated to Alexa Fluor 488, 549, or 647 (Invitrogen; Waltham, MA) and used at 1:1000. HaloTags were visualized with JaneliaFluor549 or 646-conjugated Halo Ligands (Promega; Madison, WI). Labeling of HaloTags was performed on live cells for 0.5-2 h at 30°C at a final concentration of 100 μM.

**Table 2.**
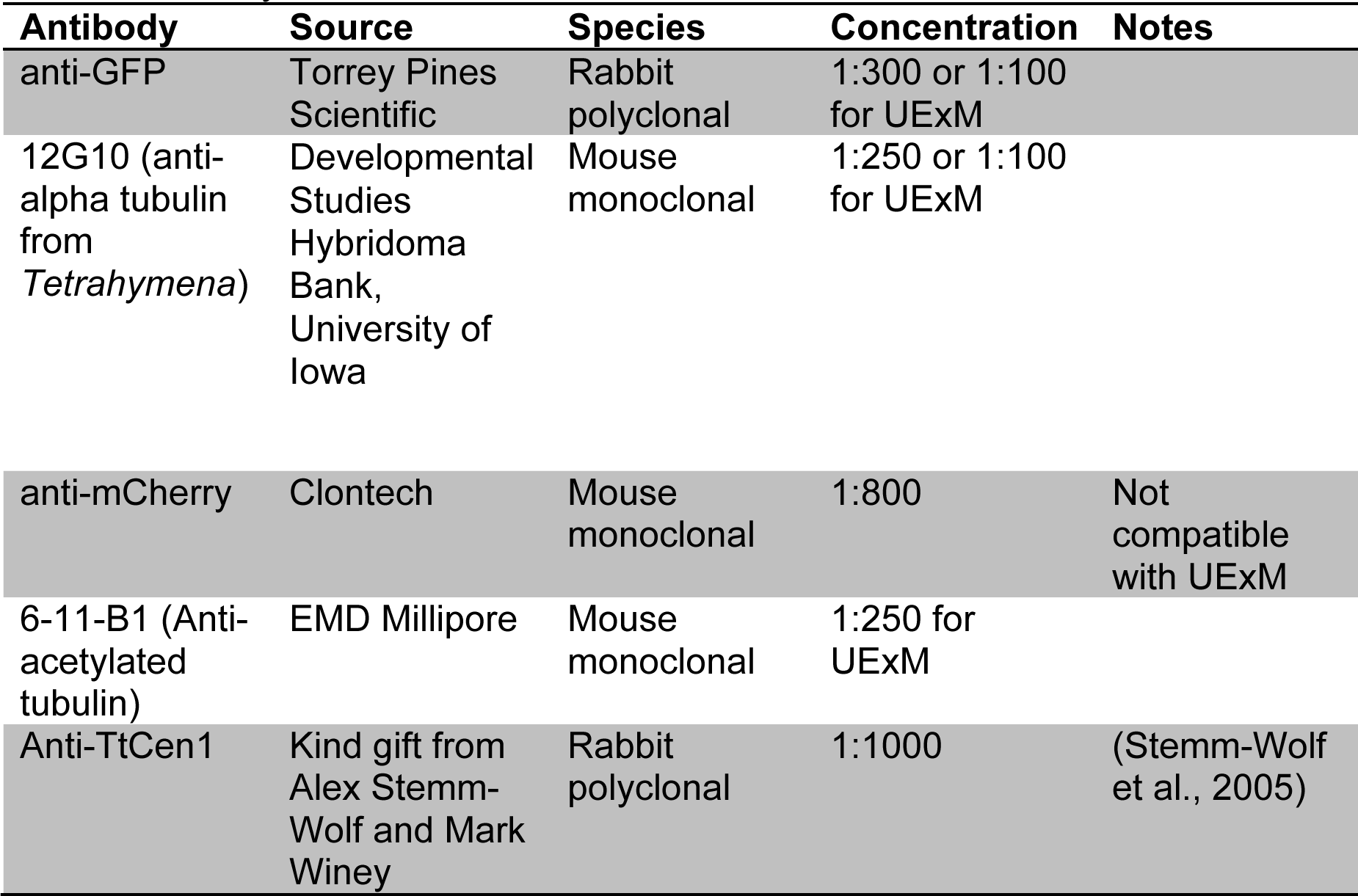
Primary Antibodies.

| **Antibody** | **Source** | **Species** | **Concentration** | **Notes** |
| --- | --- | --- | --- | --- |
| anti-GFP | Torrey Pines Scientific | Rabbit polyclonal | 1:300 or 1:100 for UExM |  |
| 12G10 (anti-alpha tubulin from *Tetrahymena*) | Developmental Studies Hybridoma Bank, University of Iowa | Mouse monoclonal | 1:250 or 1:100 for UExM |  |
| anti-mCherry | Clontech | Mouse monoclonal | 1:800 | Not compatible with UExM |
| 6-11-B1 (Anti-acetylated tubulin) | EMD Millipore | Mouse monoclonal | 1:250 for UExM |  |
| Anti-TtCen1 | Kind gift from Alex Stemm-Wolf and Mark Winey | Rabbit polyclonal | 1:1000 | (Stemm-Wolf et al., 2005) |

### Confocal Microscopy

Confocal fluorescence images were acquired using a Nikon Ti Eclipse inverted microscope with a Nikon 100x Plan-Apo objective, NA 1.45, at 23°C and Andor iXon X3 camera, and CSU-X1 (Yokogawa) spinning disk. Images were acquired using Slidebook6 or Slidebook23 imaging software and analyzed using ImageJ image analysis software (https://imagej.net). All images were acquired with exposure times between 50 and 500 ms, depending on the experiment and the channel of acquisition.

### Ultrastructure Expansion Microscopy

One milliliter of cells was pelleted at 0.5xg for 3 minutes at room temperature and then resuspended in 0.5 mL of Formaldehyde / Acrylamide solution (1.4% / 2%) for 2-5 h at 37°C with nutation. Cells were pelleted again using the same settings and 5 μl of the pellet was placed on a piece of parafilm laid flat within a humid chamber lying on ice. Thirty five microliters of monomer solution (19% Sodium Acrylate, 10% Acrylamide, 0.1% N’,N’-methylenebisacrylamide, 1x PBS)/ 0.5% APS/ 0.5% TEMED mixture was added on top of the 5 μl cell drop and an 18 x 18 mm #1.5 coverglass was laid gently on top to form a thin layer of gel between the glass and parafilm. Gels were formed on ice for 5 minutes then transferred to 37°C for 1 hour. After gelation, the coverslips were flipped over, with the gel remaining adhered to the glass. Several 4 or 6 mm diameter punches were cut into the gel using a homemade, 3D printed punch, which created multiple gel “discs” from a single sample. The gels were then transferred to 1.7 mL microcentrifuge tubes and 1.5 mL of denaturation buffer (200 mM SDS, 200 mM NaCl, 50 mM Tris in water, pH 9) was added. The tubes were incubated at 95°C for 1.5 hours. After denaturation, the gels were expanded twice in 50 mL conical vials with double distilled water, then overnight in 50 mL of ddH_2_O.

Gels were stained as follows. A single gel disc was transferred to a 12-well plastic dish and washed with 1x PBS two times for 15 minutes. PBS was removed carefully to avoid damage to the gel and 200-500 μl of primary antibody solution in 2% BSA-PBS was added to the wells and incubated for 2 h at 37°C, or overnight at 4°C, with gentle rocking. Gels were washed three times with 1 mL of PBS-Tween20 0.1%, for 10 minutes each. Washing solution was gently removed and 200-500 μl of secondary antibody solution in 2% BSA-PBS was added to the wells and incubated for 2 h at 37°C with gentle rocking. Gels were then washed twice with 1 mL of PBS-Tween20 0.1%, for 10 minutes each and re-expanded in ddH20 as described above.

For imaging, gels were sandwiched between two poly-L-lysine coated 22 x 60 mm #1.5 coverslips then clamped to the stage mount on either confocal or SIM microscopes. Expansion factors obtained in these experiments ranged from 2.5-3.75 fold, using the diameter of the BB as a molecular ruler. All scale bars in UExM images represent the scale of the acquired image and are not corrected for expansion factor.

### SIM Microscopy

SIM imaging was performed on a Nikon N-SIM system using a Ti2 inverted microscope with a 100x CF160 Apo-chromoat superresolution/TIRF NA 1.49 objective with correction collar (Nikon Instruments) and a sCMOS camera (ORCA-Flash4.0, Hamamatsu). Images were collected at 25°C and reconstructed using the slice reconstruction algorithm (NIS Elements).

### Statistical Analyses

Experiments were performed with three biological replicates. The total number of cells and / or BBs analyzed is described in the figure legends or in the Materials and Methods section. Prism6.2e (GraphPad Software) was used for graphing and statistical analysis. All data sets were tested for normality using the D’Agostino-Pearson omnibus normality test. Normally distributed data were analyzed using the Student’s t-test, whereas non-normal data were analyzed using the Mann-Whitney test. Exact P-values are reported on each graph. Error bars represent standard deviation.

**Figure S1:**
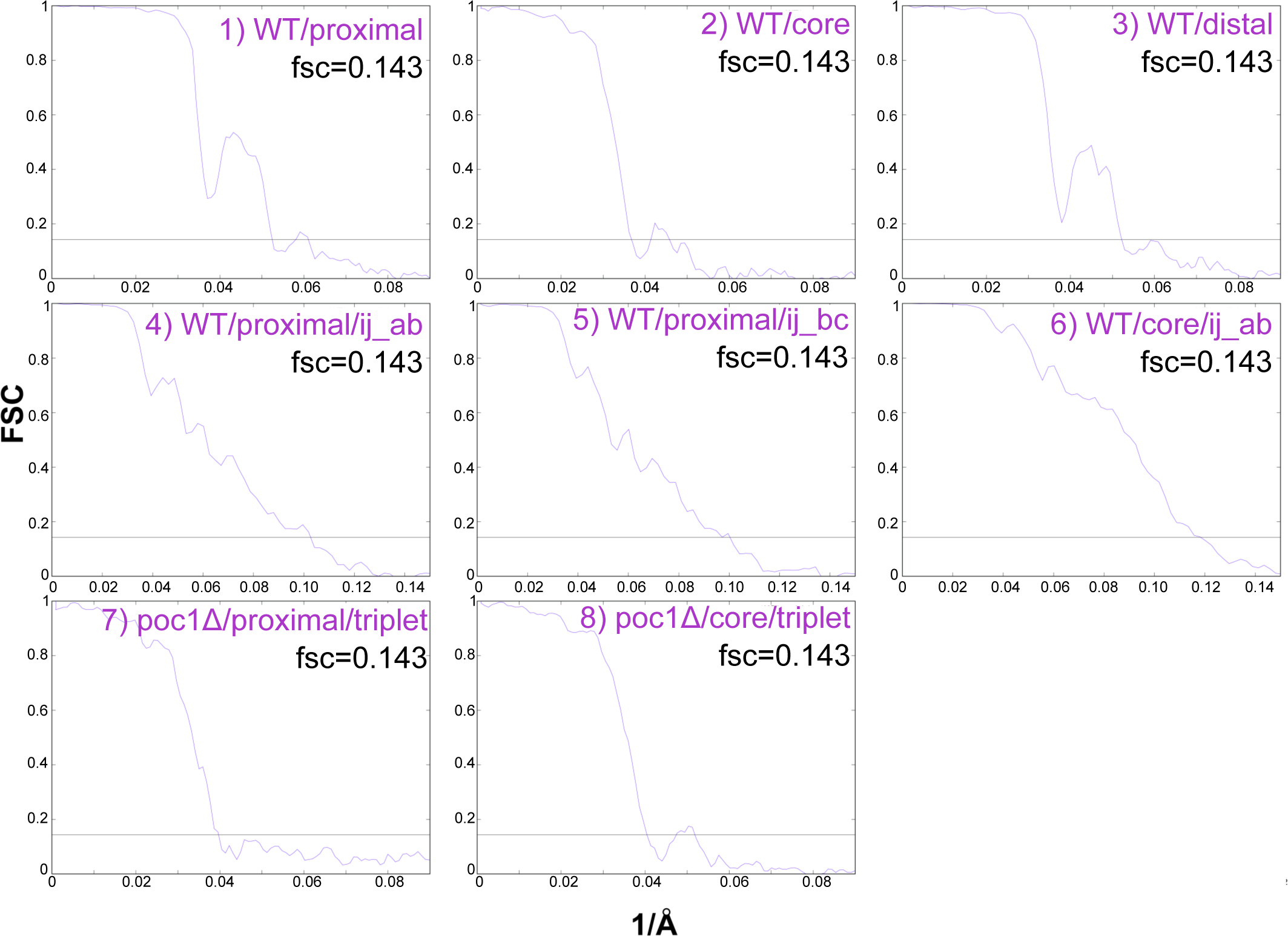
Assessing resolution of subtomogram averages by Fourier shell correlation. Plots graph the Fourier shell correlation as a function of resolution (1/Å) for each structure reported in this work (accompanies Table 1).

**Figure S2:**
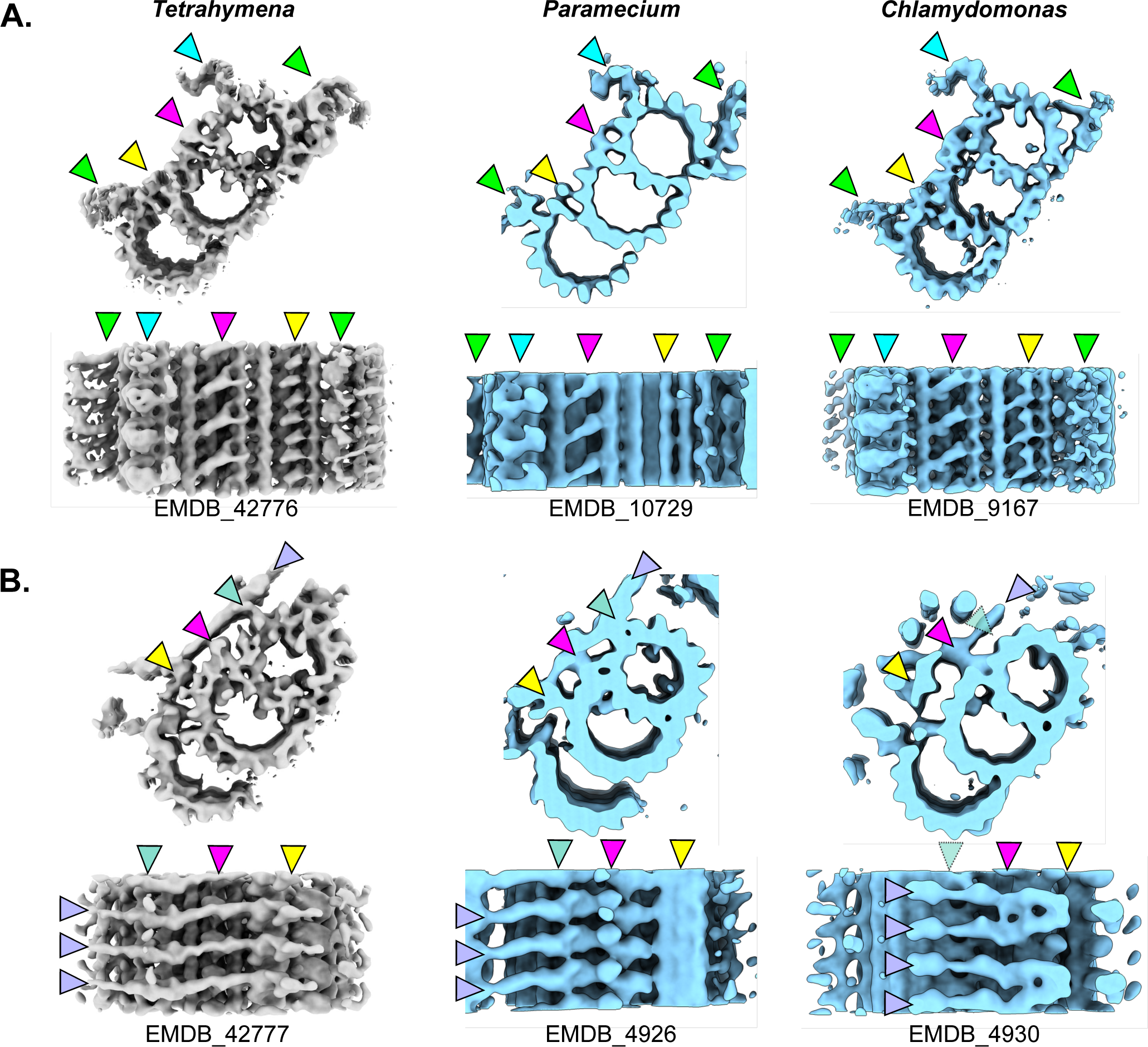
Structural features of the *Tetrahymena* BB and comparison to other species. A) Structure of TMTs from the proximal region of the BB from *Tetrahymena* (this work), *Paramecium* (EMDB_10729), and *Chlamydomonas* (EMDB_ 9167). Arrowheads denote structural features of the proximal region: A-C linkers (green), pinhead (cyan), A-B inner junction (magenta), and B-C inner junction (yellow). B) Structure of TMTs from the central core region of the BB from *Tetrahymena* (this work), *Paramecium* (EMDB_4926), and *Chlamydomonas* (EMDB_ 4930). Arrowheads denote structural features of the core region: inner scaffold (lavender), A03 attachment site (teal) A-B inner junction inner scaffold attachment site (magenta), and B-C inner junction inner scaffold attachment site (yellow). Opaque teal arrowhead with dashed outline in the *Chlamydomonas* core region structure indicates a missing A03 attachment.

**Figure S3:**
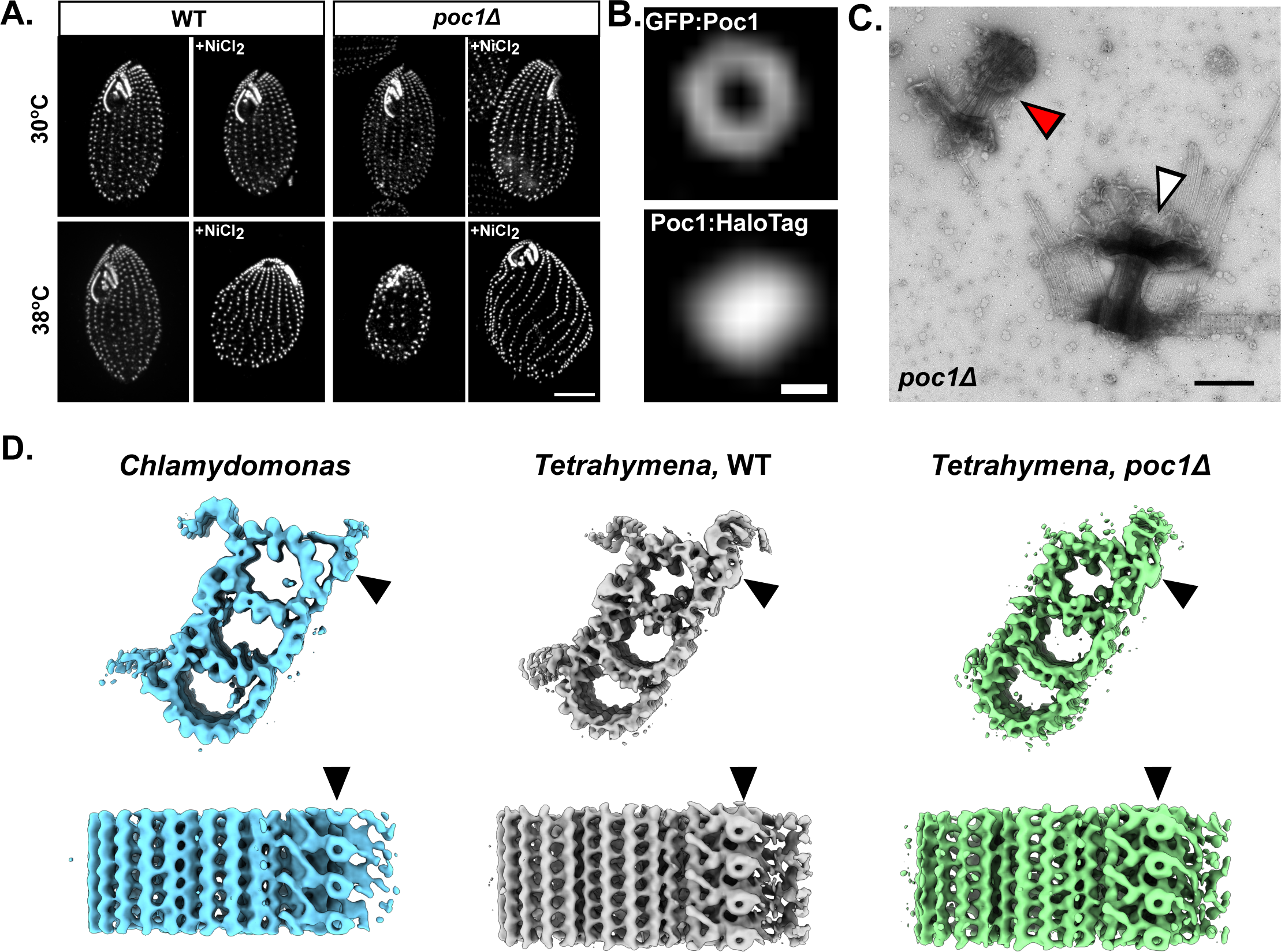
Poc1 is required to resist ciliary forces and its WD40 domain is not in the A/C linker. A) Inhibition of ciliary beating with NiCl_2_ rescues the poc1Δ BB loss phenotype at high force. This was also shown in (Meehl et al., 2016). Cells were stained with anti-centrin antibody. Scale bar is 10 μm. B) SIM images of Poc1 N-terminal tag (GFP) versus C-terminal tag (HaloTag). Rings are only seen with N-terminal tags. Scale bar is 100 nm. C) Negative stain of isolated poc1Δ BBs demonstrating that some BBs maintained their overall structure (white arrowhead), whereas some were severely disintegrated (red arrowhead). Scale bar is 400 nm. D) The doughnut shaped density previously proposed to be Poc1 in the A/C linker of *Chlamydomonas* BBs is also present in WT *Tetrahymena* BBs and remains present in poc1Δ *Tetrahymena* BBs. Proximal end TMTs from *Chlamydomonas* (left, blue), WT *Tetrahymena* (middle, gray), and poc1Δ *Tetrahymena* BBs (right, green) in cross section (top) and exterior side views (bottom). Arrowheads point to the location of the density previously proposed to be Poc1 (Li et al., 2019).

**Figure S4:**
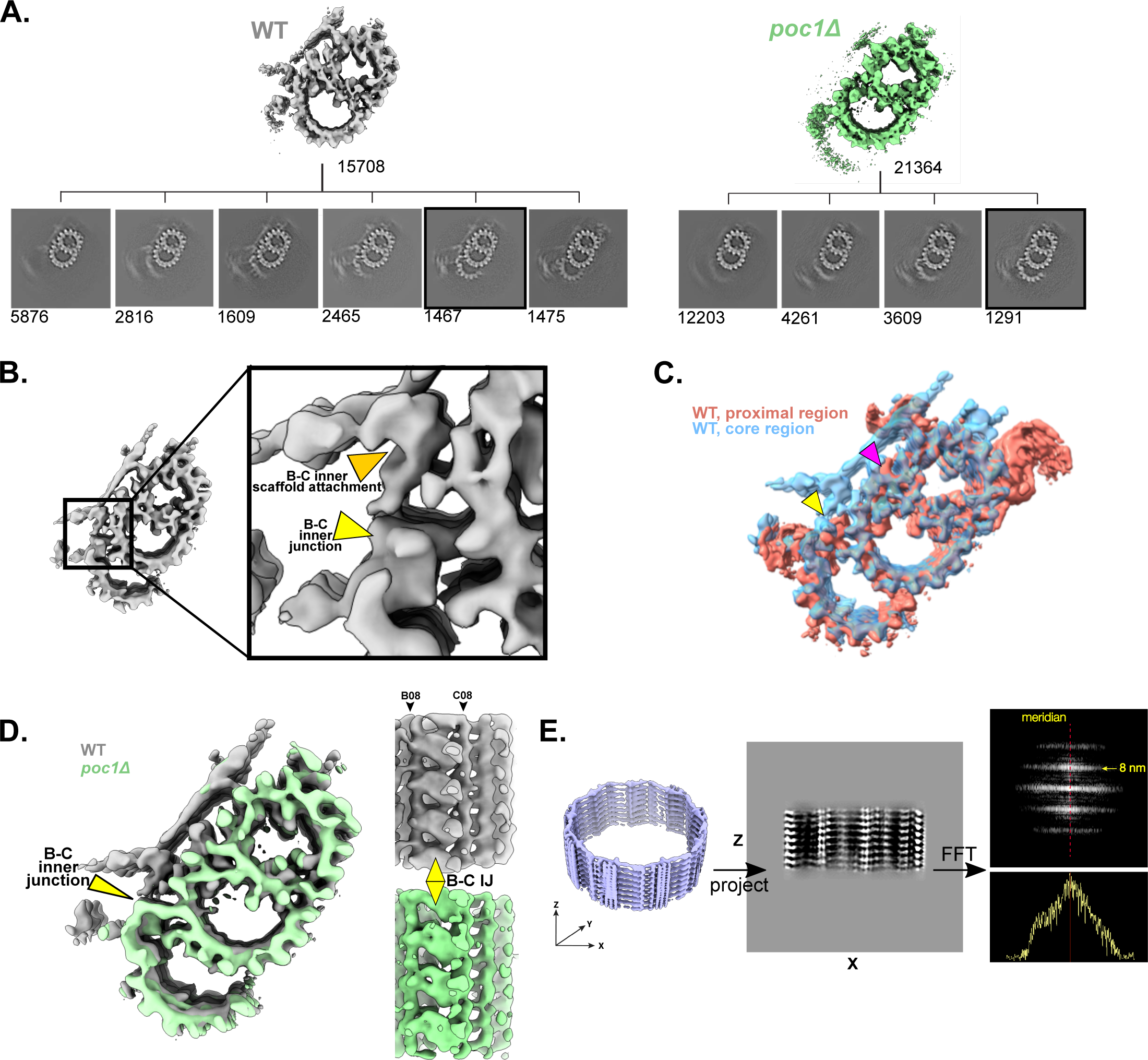
Classification of TMT from WT and *poc1Δ*, the BC inner junction in the core region does not contain Poc1, and the inner scaffold is a stack of circular rings in *Tetrahymena*. A) Subtomogram averages and classifications of the WT and *poc1Δ* TMT from the core region. Note the loss of C-tubules in WT TMTs, indicating C-tubules are labile in the core region of the BB. The numbers of subtomograms in each class is indicated. Classes with black boxes around them indicate the complete TMT structure from WT and *poc1Δ*. B) Cross-sectional view of the complete TMT from the WT core region, zoomed in on the B-C inner junction. Arrowheads point to the B-C inner junction structure (yellow) and the structure of the B-C inner junction attachment to the inner scaffold (marigold). C) Comparison of WT *Tetrahymena* TMTs from the proximal (salmon) and core (blue) regions. The overlay reveals the B-C inner junction is different in architecture between the proximal and core regions. D) Overlay of complete WT (gray) and poc1Δ (green) TMTs from the core region in cross-section (left) and luminal side view (right). Although the B-C inner junction inner scaffold attachment site is lost in *poc1Δ*, the structure of the B-C inner junction itself is not different between WT and *poc1Δ*. Also note that the C-tubule links to the B-tubule using apparently non-tubulin structures that are retained in complete poc1Δ TMTs. E) The *Tetrahymena* inner scaffold is a series of stacked rings. Left, the structure of the inner scaffold backfit to the BB core region. Middle, projection of the density onto the X-Z plane. Right, Fourier transform of the projected density (top) and a linescan of intensity at 8 nm layer line (bottom) indicates its Bessel order close to zero, suggesting the scaffold is a zero-start helix. See methods for a description of this analysis.

**Figure S5:**
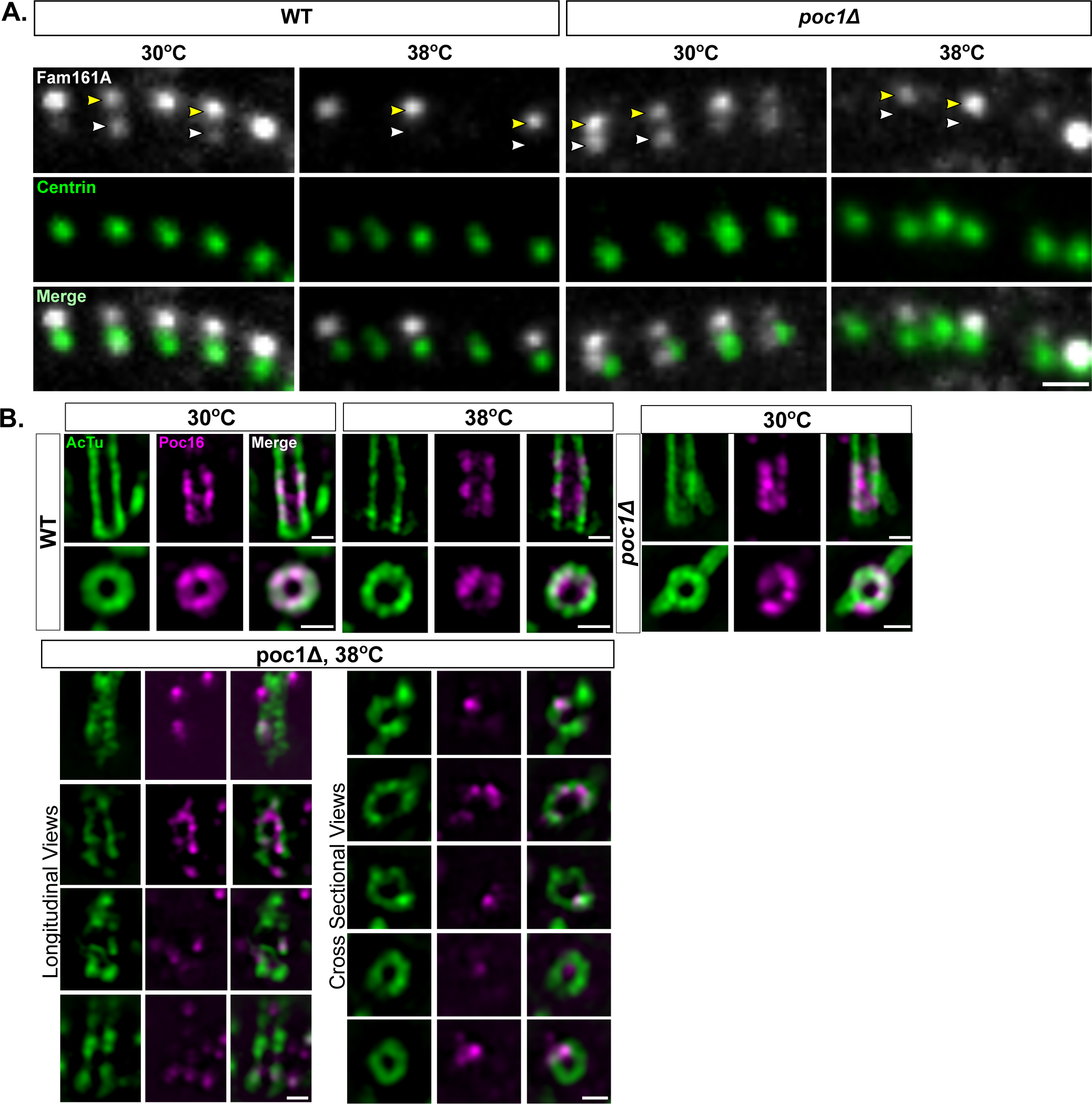
Fam161A is lost at the core region in WT and poc1Δ cells, and BB defects in poc1Δ at high force are associated with loss of Poc16. A) Side view of a ciliary row from WT (left) and poc1Δ (right) cells expressing Fam161A:mCherry at 30°C and 38°C. The cell exterior is at the top of the images. Cells were co-stained with anti-mCherry and anti-centrin antibodies. Fam161A has two longitudinal localizations, core (white arrowhead) and presumed transition zone (yellow arrowhead), with the latter population being the most consistent in each BB in low force conditions. Upon shift to 38°C, Fam161A is lost preferentially from the core region in both WT and poc1Δ cells. Scale bar is 1 μm. B) UExM-SIM images of WT and poc1Δ cells at 30°C and 38°C expressing Poc16:GFP. Representative longitudinal and cross sectional images are shown for WT at both temperatures and poc1Δ at 30°C. For poc1Δ at 38°C a montage of four longitudinal views (bottom left) and five cross sectional views (bottom right) are shown. Figure accompanies Fig. 6F. Scale bars are 500 nm.

## Abbreviations used

BB: basal body
TMT: triplet microtubule
DMT: doublet microtubule
UExM: Ultrastructural Expansion Microscopy
cryoET: electron cryo-tomography

## ACKNOWLEDGEMENTS

The authors would like to thank Mark Winey for helpful conversations and partial funding of Sam Li during the course of the work. In addition, the University of Colorado School of Medicine Cryo Electron Microscopy shared resource center, especially Brian Wimberly, Dave Farrell, Peter Van Blerkom, and Eduardo Romero Camacho for help and expertise in preparing and screening EM grids. We also thank Jose-Jesus Fernandez (CINN-CSIC, Spain) for advice on cryoET image processing, David Bulkley and Eric Tse (UCSF) for assistance with cryoET data collection, Tom Goddard (UCSF) for advice on AlphaFold and ChimeraX, and UCSF Wynton HPC for computation support during this study.

This research was funded by NIH GM140813 (C.G. Pearson), NIH R35GM118099 (D.A. Agard), NIH 2R01GM127571-05A1 (M. Winey) and NIH-NIGMS 5F32GM122239 (M.D. Ruehle).

## AUTHOR CONTRIBUTIONS

M.D. Ruehle conceived and performed experiments, prepared BBs and cryoET grids, performed data analysis, created figures, and wrote and revised the manuscript. S. Li collected cryo-tomography data, performed data analysis for structure determination, created figures, wrote sections of the manuscript describing the pseudoatomic models, and revised the manuscript. D.A. Agard and C.G. Pearson conceived of the cryo-tomography experiments, supervised, revised the manuscript, and provided resources and funding for the project.

